# Multi-State Modeling of G-protein Coupled Receptors at Experimental Accuracy

**DOI:** 10.1101/2021.11.26.470086

**Authors:** Lim Heo, Michael Feig

**Author notes:** Corresponding author: 603 Wilson Road, Room 218 BCH, East Lansing, MI 48824, USA.

## Abstract

The family of G-protein coupled receptors (GPCRs) is one of the largest protein families in the human genome. GPCRs transduct chemical signals from extracellular to intracellular regions via a conformational switch between active and inactive states upon ligand binding. While experimental structures of GPCRs remain limited, high-accuracy computational predictions are now possible with AlphaFold2. However, AlphaFold2 only predicts one state and is biased towards either the active or inactive conformation depending on the GPCR class. Here, a multi-state prediction protocol is introduced that extends AlphaFold2 to predict either active or inactive states at very high accuracy using state-annotated templated GPCR databases. The predicted models accurately capture the main structural changes upon activation of the GPCR at the atomic level. For most of the benchmarked GPCRs (10 out of 15), models in the active and inactive states were closer to their corresponding activation state structures. Median RMSDs of the transmembrane regions were 1.12 Å and 1.41 Å for the active and inactive state models, respectively. The models were more suitable for protein-ligand docking than the original AlphaFold2 models and template-based models. Finally, our prediction protocol predicted accurate GPCR structures and GPCR-peptide complex structures in GPCR Dock 2021, a blind GPCR-ligand complex modeling competition. We expect that high accuracy GPCR models in both activation states will promote understanding in GPCR activation mechanisms and drug discovery for GPCRs. At the time, the new protocol paves the way towards capturing the dynamics of proteins at high-accuracy via machine-learning methods.

## INTRODUCTION

High-resolution protein structures provide detailed mechanistic information at the atomistic level about biological function and serve as starting points for structure-based drug design to develop small molecules that control protein behavior.^1,2^ Such high-resolution structures have been acquired by experimental methods such as X-ray crystallography, nuclear magnetic resonance spectroscopy, and cryo-electron microscopy. An increasingly powerful alternative is computational protein structure prediction. Predictions based on homology templates have long provided confident models when there is an experimentally determined structure for a close homolog.^3,4^ More recently, protein structure prediction based on machine learning methods such as AlphaFold2 (AF2)^5^ and RoseTTAFold^6^ has also become able to generate accurate models for essentially any sequence, even when homologs are not available. These methods mainly rely on neural network models that have learned how to deduce inter-residue relationships based on co-evolutionary couplings together with high-resolution structure generation modules trained on known experimental structures. The accuracy of the resulting models may approach experimental accuracy, and at least for some applications, the computational models may be sufficient for further studies. However, limitations remain with respect to capturing structural dynamics that can lead to multiple conformations. Proteins often possess multiple conformational states to perform their biological roles, whereas prediction methods are generally trained to predict a single, native state for a given sequence. As may be expected, models produced by AF2 during CASP14 (14^th^ critical assessment of protein structure prediction) typically varied little, although different models for one target could match multiple conformational states accurately.^7,8^ Given the overall success of AF2 in producing high-accuracy models, the question of whether and how multiple states can be predicted for a given protein needs further investigation and is the subject of this study.

G-protein coupled receptors (GPCRs) comprise one of the largest protein families in the human genome.^9^ They are involved in various biological roles such as behavior regulation and regulation of immune system activity.^1,2,10,11^ They are subdivided into several classes according to sequence homology and functional similarity: class A (rhodopsin), B1 (secretin), B2 (adhesion), C (glutamate), F (frizzled), T (taste 2).^12^ Dysregulation of GPCRs can cause diseases including cardiovascular disease^11^ and Parkinson’s disease.^10^ GPCRs are attractive targets for drug discovery, and indeed, around one third of drugs approved by the U. S. Food and Drug Administration (FDA) target GPCRs.^1,13^ GPCRs are integral membrane proteins with a common topology that consists of seven transmembrane helices. GPCRs generally function by transmitting chemical signals from the extracellular to the intracellular region. The transmission is usually triggered by the binding of agonists on the extracellular side. The binding alters the conformation of a GPCR from the inactive to an active state. The active state can then be sensed on the intracellular side via binding of a transducer molecule, an active G-protein or a β-arrestin. The details of the activation mechanism vary depending on the class of the GPCR and they are still under investigation via experimental and computational approaches.^14–20^

Experimental structures of GPCRs are available since the first GPCR structure, for bovine rhodopsin, was resolved in 2000.^21^ The number of experimental GPCR structures has steadily increased since.^2^ As of January 05, 2022, 689 GPCR structures have been deposited to the protein data bank (PDB).^22^ However, a majority of GPCR structures are still unknown. Known structures comprise only 112 out of 401 human non-olfactory receptors. Moreover, even although GPCRs can have multiple conformational states, only 74 and 65 GPCRs were determined in active and inactive states, respectively, and only 36 GPCRs have experimental structures in both activation states. The lack of high-resolution GPCR structures has made it difficult to understand GPCR activation mechanisms and enable structure-based drug design for GPCRs. Computational protein structure prediction methods targeted at GPCRs have been used to generated models where experimental structures are not available. Among them, GPCR-I-TASSER^23^, RosettaGPCR^24^, GPCRdb^25,26^, and RoseTTAFold^6^ have resulted in structure databases of the whole GPCR proteome via computational modeling. The Zhang group designed a specialized version of I-TASSER^27^ for GPCRs.^23^ It was based on using GPCR structure-specific features with I-TASSER’s template-based and - free modeling pipeline. RosettaGPCR performed template-based modeling using Rosetta to predict GPCR structures in the inactive state.^24^ GPCRdb maintains multiple databases related to GPCRs including experimental structures of GPCR with activation state annotations.^25,26^ GPCRdb also contains predictions for GPCRs in active and inactive forms via homology modeling by using templates in the corresponding states. However, the accuracy of these GPCR models has not been rigorously benchmarked, especially with respect to reproducing differences between active and inactive states and in light of the recent advances in structure prediction accuracy.^25^ Baek *et al*. demonstrated that RoseTTAFold can model GPCRs in the active and inactive states by providing activation-state specific templates.^6^ The method generated reasonable models for both states. However, generating very accurate active state models was less likely unless there were templates with high sequence identities, thus, there was still room for improvements.

Here, we focus on using AF2 for modeling GPCRs, especially with respect to accurately predicting both active and inactive states for a given GPCR. As described below in more detail, the regular AF2 protocol can predict accurate GPCR models, more accurate than template-based modeling, but it was not possible to predict models in multiple states for most of the benchmarked GPCRs. The predicted models were in only one GPCR state with a preference for inactive states. To overcome this limitation, we describe here a modified protocol that enables the modeling of multiple states via AF2. Briefly, this was accomplished by using state-annotated structure databases together with state specific structural templates and a modification of the MSA input features. While the approach is general in principle, we applied it here to generate accurate models of both, active and inactive states of a given GPCRs. The protocol was tested for GPCRs where experimental structures are available and the resulting GPCR models were further examined in the context of predicting ligand binding poses. The new protocol for multi-state modeling via AF2 and results are described in more detail in the following.

## RESULTS AND DISCUSSION

### Modeling of GPCRs using AF2

The original AF2 showed outstanding performance in protein structure prediction during CASP14^5,7,28,29^ where target proteins were mostly globular soluble proteins but also included three transmembrane proteins.^8^ As benchmarked here, we find that GPCRs, one of the major transmembrane protein families, can also be predicted accurately using AF2. For recently experimentally determined GPCR structures that were not included during AF2 training, 34 out of 68 (50%) GPCRs were predicted at an accuracy of better than 1.5 Å accuracy in terms of Cα-RMSD (root-mean-square-deviation) for the transmembrane helices (TM-RMSD) with respect to any experimental structures regardless of their activation state; the median accuracy was 1.49 Å. (**Figure S1**) For comparison, those GPCR structures were also modeled via template-based modeling (TBM) with the same structural templates from the PDB70 database^30^ used in the AF2 predictions. As expected, the resulting models were less satisfactory than the AF2 predictions. Only 3 out of 68 (4%) structures had RMSD values below 1.5 Å for the TM domain, and the median TM-RMSD was 2.71 Å.

AF2 only modeled either the active or inactive state for a given GPCR sequence, though, not both. Interestingly, when its performance was analyzed on human GPCRs that were experimentally determined in both states, the top-ranked models were more likely to be in the inactive state rather than the active state for all GPCR classes except for class B1. (**Figures S2–4 and Table S1**) Out of 15 multi-state human GPCRs, 11 models were closer to their inactive state experimental structures than to their active structures. As a consequence, when comparing AF2 models strictly against the experimental structures, AF2 predictions were highly accurate for the inactive structures with a median TM-RMSD of 1.33 Å while 19 out of 30 (63%) GPCRs were predicted within 1.5 Å in TM-RMSD. In contrast, the AF2 models were dissimilar to active state GPCR structures with a median TM-RMSD of 1.86 Å. For comparison, the median TM-RMSD between active and inactive state structures was 2.26 Å. This bias may originate from the larger number of experimental GPCR structures in inactive states (178) than active states (60) that were available for training.

When multiple output models with different AF2 network model parameters were generated, predictions for most GPCRs remained very similar (**Figure S5**). However, for some cases, the resulting models varied more. For example, AF2 predictions for the human parathyroid hormone receptor (human PTH1R; UniProt ID Q03431) generated a diverse ensemble of structures. While the top-ranked model was in the active state with a TM-RMSD of 1.65 and 4.72 Å with respect to experimental structures in the active and inactive states, lower-ranked models were in the middle of the active and inactive states. The fifth-ranked model had structural similarities of 2.46 and 2.66 Å in TM-RMSD with respect to the active and inactive structures. Based on this analysis, we inferred that although AF2 is not designed *per se* to generate ensembles consisting of multiple, functionally relevant states, the AF2 neural network model can in principle predict them.

Using either multiple sequence alignments (MSA) or structural templates as input features was enough for high accuracy structure prediction using AF2, based on an ablation study. (**Figure S1**-**3**). When we excluded either MSA or structural template-based features, there was little loss in accuracy when structural templates were not used. The median TM-RMSD was 1.55 Å for recently determined human GPCRs. *vs*.1.49 Å using the original AlphaFold protocol. This finding is consistent with a previous analysis of AF2 for targets where sufficiently deep MSAs are available.^5^ The proteins in our benchmark sets have at least thousands of homologous sequences in their MSA. On the other hand, when using input features from templates, but without MSAs - essentially this is “template-based modeling” via AlphaFold - it was still possible to predict overall high accuracy models. The median TM-RMSD was 1.84 Å, better than template-based models with the same templates using MODELLER^31^ with a median TM-RMSD of 2.71 Å. (**Figures S1–3**). Based on these tests, we found out that AF2 is capable of accurate modeling with a subset of input features or less accurate structural inputs. However, a bias towards more accurate modeling of inactive states vs. active states remained.

### Multi-state modeling of GPCRs

Activation state-annotated GPCR databases could be simply used instead of the standard template database, PDB70, to model GPCRs in a desired functional state. However, using such curated template databases was not sufficient to make meaningful changes. (**Figures S2-4**) For all the multi-state GPCRs, there was little change from the result with the standard template database. This suggests that simply using templates according to a conformational state as input to AF2 has little impact when deep MSAs were provided as inputs.

On the other hand, when MSA input features were removed and state-annotated GPCR databases were used, it became possible to generate highly accurate models for both active and inactive state structures of GPCRs (**Figure 1** and **Figures S1–3, 6**). In this manner, the activation state-annotated structural template databases could be used to guide the AF2 modeling network towards a specific activation state based on homology, but without interference from MSA-based contacts that are still biased by conformational state preferences according to training. This strategy led to greatly improved active state predictions while maintaining the previously achieved high accuracy for inactive state structures (**Figure 1a**). Many of the GPCRs (35 out of 49, 71%) were predicted within 1.5 Å TM-RMSD, and the median of the active state modeling accuracy in TM-RMSD was 1.12 Å. For inactive states, structures were predicted with a median TM-RMSD of 1.41 Å, and 17 out of 30 models predicted at high-accuracy (*i.e*., < 1.5 Å TM-RMSD) (*c.f*, 19 high-accuracy models and 1.33 Å for the original AF2). The performance difference between two methods was evaluated using a paired *t*-test with the null hypothesis that two methods had identical performance. We found that differences were statistically significant only for the active state predictions according to p-values for the active and inactive states of 8.2×10^-8^ and 0.37, respectively. Moreover, for 10 targets among the 15 multi-state GPCRs, our protocol successfully modeled the correct activation states based on the active state model being closer to the active state experimental structures than to the inactive experimental structure and *vice versa* for the inactive state model.

**Figure 1.**
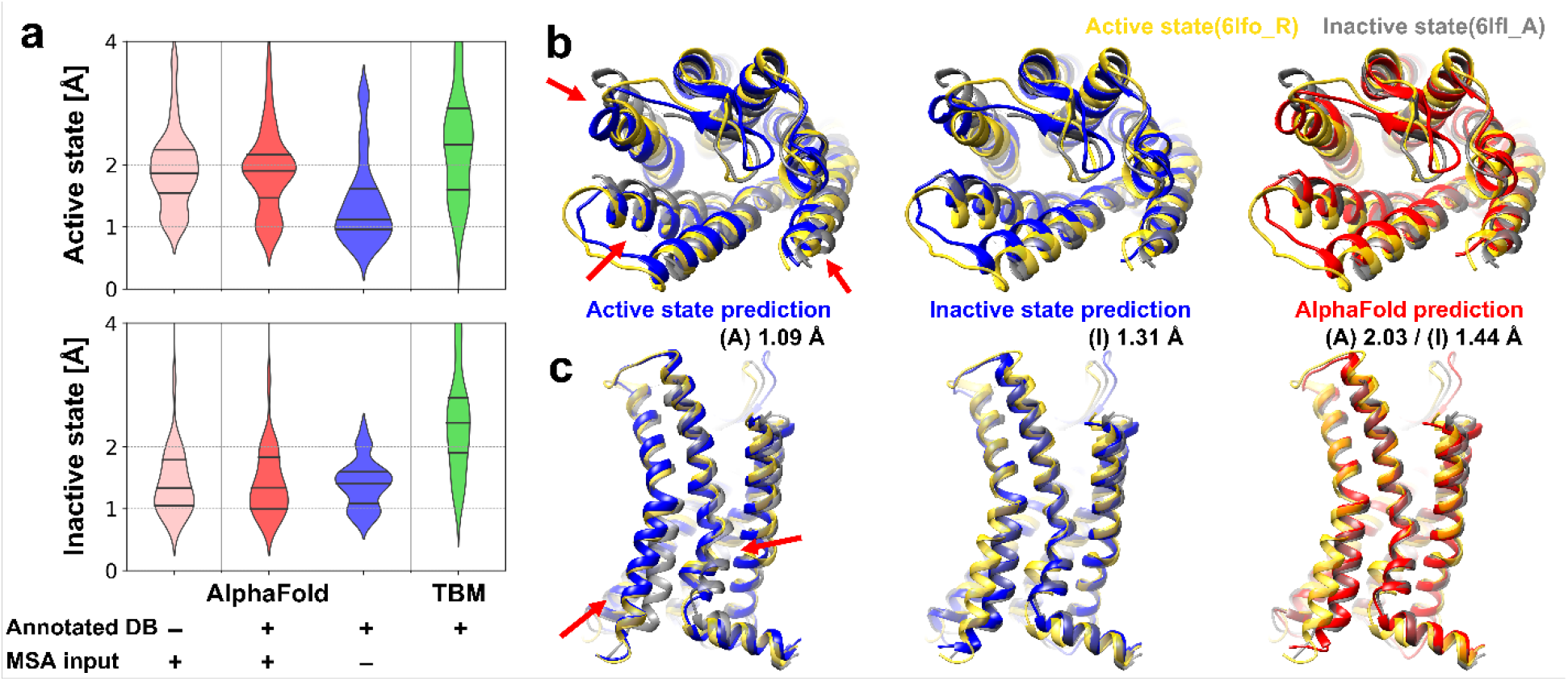
Modeling of active and inactive states of human G-protein coupled receptor structure (GPCR). (a) Modeling accuracies for human GPCRs that have both active and inactive state experimental structures using various modeling protocols based on AlphaFold and template-based modeling (TBM). Cα-RMSDs are measured with respect to active and inactive forms of transmembrane helices. (TM-RMSD) Distributions of modeling accuracies are shown as violin plots with black lines indicating three quartiles. (b and c) Multi-state modeling of human C-X-C chemokine receptor type 2 (UniProt ID P25025, CXCR2_HUMAN). (b) View from the extracellular region; (c) Side-view. Models predicted as active and inactive forms using customized GPCR databases via AF2 without MSA input features are shown in blue. A predicted model using AF2 with MSA input features and the standard PDB70 database is shown in red. Experimental structures of active state (yellow, PDB ID 6lfo_R) and inactive state (gray, PDB ID 6lfl_A) are compared to the predictions. Selected conformational differences between states are highlighted by red arrows.

Furthermore, we analyzed the modeling performance as a function of the GPCR class. (**Table S1**) The multi-state modeling protocol was successful for predicting class A GPCRs in both active and inactive states. For class F GPCRs, it was satisfactory as well, however, there were only three targets in the benchmark set. It also generated high-accuracy models for class B1 GPCRs in the active state. Model qualities of inactive state class B1 GPCRs were slightly less satisfactory than the active state models, but they were still better than the original AF2 models. Similarly, for class C, modeling was successful for the inactive states but less successful for the active state. A poor model was generated for a class B2 GPCR, but based on only one example we cannot draw more general conclusions about modeling class B2 GPCRs. On the other hand, the original AF2 protocol showed biases towards a certain activation state depending on the GPCR class. It showed strong preferences for the inactive state for class A, C, and F, while it generated active state conformations for class B1 GPCRs. For those classes for which the original AF2 had biases toward the inactive state, models in the state by our multi-state protocol were comparable to the AF2 models (class A) or slightly worse than AF2 (class C and F). In the AF2 training set, there were both active and inactive conformations for class A, while there were only inactive state structures for classes C and F. Because of these, AF2 may have stronger preference in the inactive state for classes C and F than class A, and as a result, it showed much better accuracy for these classes. Since we sacrificed information that can be inferred from the MSA for modeling both states, it was less likely that we could predict better models for these states, and thus we could obtain similar performance to AF2, at best. For class B1, our protocol predicted more accurate active state models than AF2 even although AF2 was likely to predict the active state for this class. Regardless of the GPCR class, our multi-state modeling protocol produced better models than AF2 for states that were not favored by the AF2.

We further assessed whether detailed aspects of structural changes upon activation of the GPCR could be captured with the multi-state modeling protocol. For example, rearrangements of TM helices during activation for the human C-X-C chemokine receptor type 2 (UniProt ID P25025, CXCR2_HUMAN) were accurately described. On the extracellular side, the movement of TM1, TM5, and TM6 when transitioning between active and inactive states was successfully modeled. (red arrow in **Figure 1b**) Also, structural changes of TM6 on the intracellular side, which enables G-protein binding, was described very accurately. (red arrow in **Figure 1c**). Beyond TM helix rearrangements, other detailed structural features correlated with the activation mechanism could be captured as well. In the example of the same protein, the changes during activation were captured very accurately up to atomistic detail in the predicted active and inactive states when compared to the experimental structures (**Figure 2**). The CXCR2 receptor has a unique activation initiation process. Movement of TM5 helix induces rearrangement of the PIF motif (P^5×50^, I^3×40^, F^6×44^) and W^6×48^. (**Figure 2a and b**)^32^ Our active and inactive state models for the receptor captured the change very accurately. In addition, our models described common structural changes for the class A GPCR correctly for the NPxxY motif (N^7×49^, P^7×50^, and Y^7×53^) and the DRY motif (D^3×49^, R^3×50^, and Y^3×51^).^19,32^ (**Figure 2c**)

**Figure 2.**
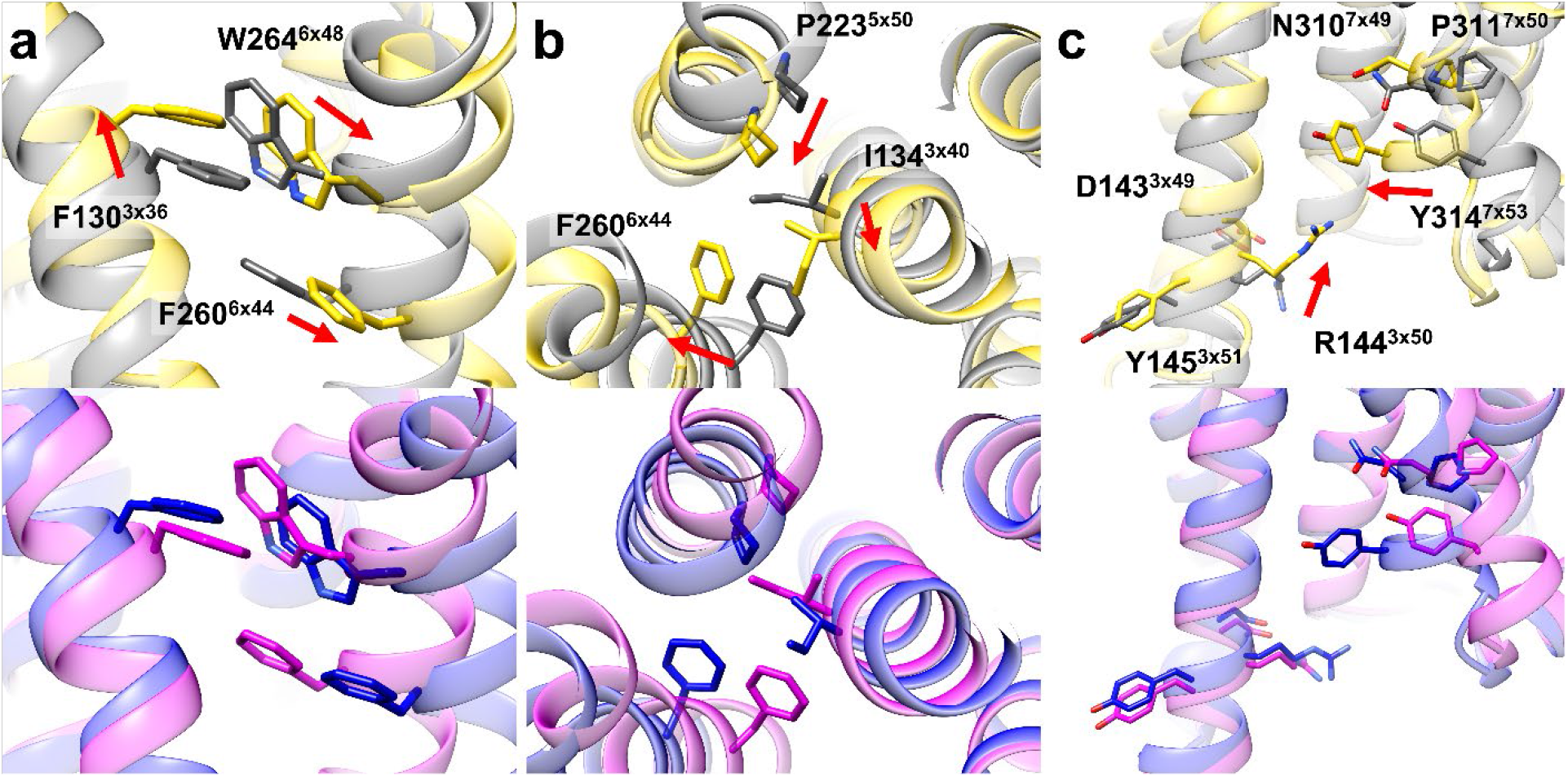
Detailed structural changes upon activation of human C-X-C chemokine receptor type 2 (UniProt ID P25025, CXCR2_HUMAN). Experimentally determined active (PDB ID 6lfo_R) and inactive (PDB ID 6lfl_A) state structures are shown in yellow and grey, respectively. Predicted active and inactive state model structures are shown in blue and magenta, respectively. Key sidechains for the activation mechanisms are shown as sticks with their residue numbers and the GPCRdb numberings, and the direction of changes is indicated by red arrows. CXCR2-specific rearrangement of (a) W^6×48^ and (b) the PIF motif (P^5×50^, I^3×40^, F^6×44^) induced by movement of the TM5 helix. (c) Common structural changes for the class A GPCRs at the NPxxY motif and DRY motif.

In addition to the transmembrane bound regions, modeling accuracies of three intra- and three extra-cellular loops (ICLs and ECLs) were evaluated. (**Figure S8**) Overall, loop structures predicted by the multi-state modeling protocol were accurate. The accuracy was mostly comparable to the original AF2 except for a few examples especially for very long ECL2s. Those unsatisfactory loop modeling mostly occurred when the sequence identity between the target protein and templates were low and loop structures were very dissimilar. As discussed above, AF2 modeling network can generate accurate models from roughly accurate input features. Thus, moderately accurate loop structure information from templates could be refined by the network as for transmembrane helices. However, substantially incorrect loop structures in the input features could not be rescued and remained incorrect.

The model accuracy with the tested modeling protocols was compared with other methods, GPCRdb^25^ and RoseTTAFold^6^. (**Figure S9**) To be comparable with our benchmark test, GPCRdb models were retrieved from an archive (2018 April) of the method’s model database. Among the benchmark set, there were 41 and 22 common GPCRs for the active and the inactive states, respectively. For the comparison with RoseTTAFold, we ran RoseTTAFold according to its multi-state modeling protocol with the same templates that we used for our protocol. As we ran the protocol by ourselves, there were the same number of targets for the comparison. Regardless of the activation state, models predicted by the multi-state modeling protocol were better than the compared methods. In comparison with RoseTTAFold, our protocol performed especially well for active state modeling. RoseTTAFold generated high-accuracy models for only 2 out of 49 targets with a median TM-RMSD of 2.16 Å. (*c.f*. 35 high-accuracy models and 1.12 Å for our protocol) AF2 generated better inactive state conformations than GPCRdb and RoseTTAFold. There were two poor predictions, and they were class B1 GPCRs that were predicted in the active state. On the other hand, for the active state, AF2 performed marginally better than the compared methods as it tended to predict in the inactive state for most of the GPCR classes.

Finally, we also predicted all human non-olfactory GPCRs in active and inactive states using our multi-state modeling protocol. We could evaluate the model accuracy for GPCR models that have experimentally determined structures as a function of the predicted local distance difference test score (pLDDT) by the AF2 network model and the maximum sequence identity of the used templates. (**Figure 3**) There was a clear relationship between the accuracy and the pLDDT as presented in Jumper *et al*.^5^ For a model that had a pLDDT higher than 90, it was likely to be accurate; 83% (29 out of 35) and 74% (14 out of 19) of the active and inactive state models had less than 1.5 Å in TM-RMSD, respectively. Among the GPCRs without experimental structures for each state, 72% (209 out of 289) and 64% (189 out of 289) of the active and inactive models are expected to be accurate as they had predicted pLDDTs higher than 90. In contrast, the pLDDT for models by original AF2 was less informative for the simultaneous assessment of multiple states. (**Figure S10**) Presumably this is because these models were similar to one of the states but not the other one. Model accuracy is also related to the maximum template sequence identity. We observed that modeling with an active state template that has a sequence identity higher than 20% usually resulted in a high quality model; 83% (30 out of 36) of the active state models had less than 1.5 Å in TM-RMSD. The metric has been widely used to assess what is likely successful template-based modeling. For example, GPCR models predicted using templates with sequence identities higher than 20–40% were assumed to be confident.^24,33,34^ Likewise, it may be used to infer the success of modeling prior to the active state modeling via our multi-state modeling protocol, and the sequence identity criteria may be lowered because the AF2 network can model accurately even with approximate information. The structural template dependence was introduced here because information from structural templates was one of the most important input features for our protocol as MSA information was discarded. For the inactive state models, we could not conclude that there is a clear template similarity dependence since there were not enough GPCRs that had low template homology. We note, that there was no such template-accuracy relationship for the original AF2 protocol. The reason is likely that structural templates have little effect when there are enough sequences in the input MSA as shown in the ablation study.

**Figure 3.**
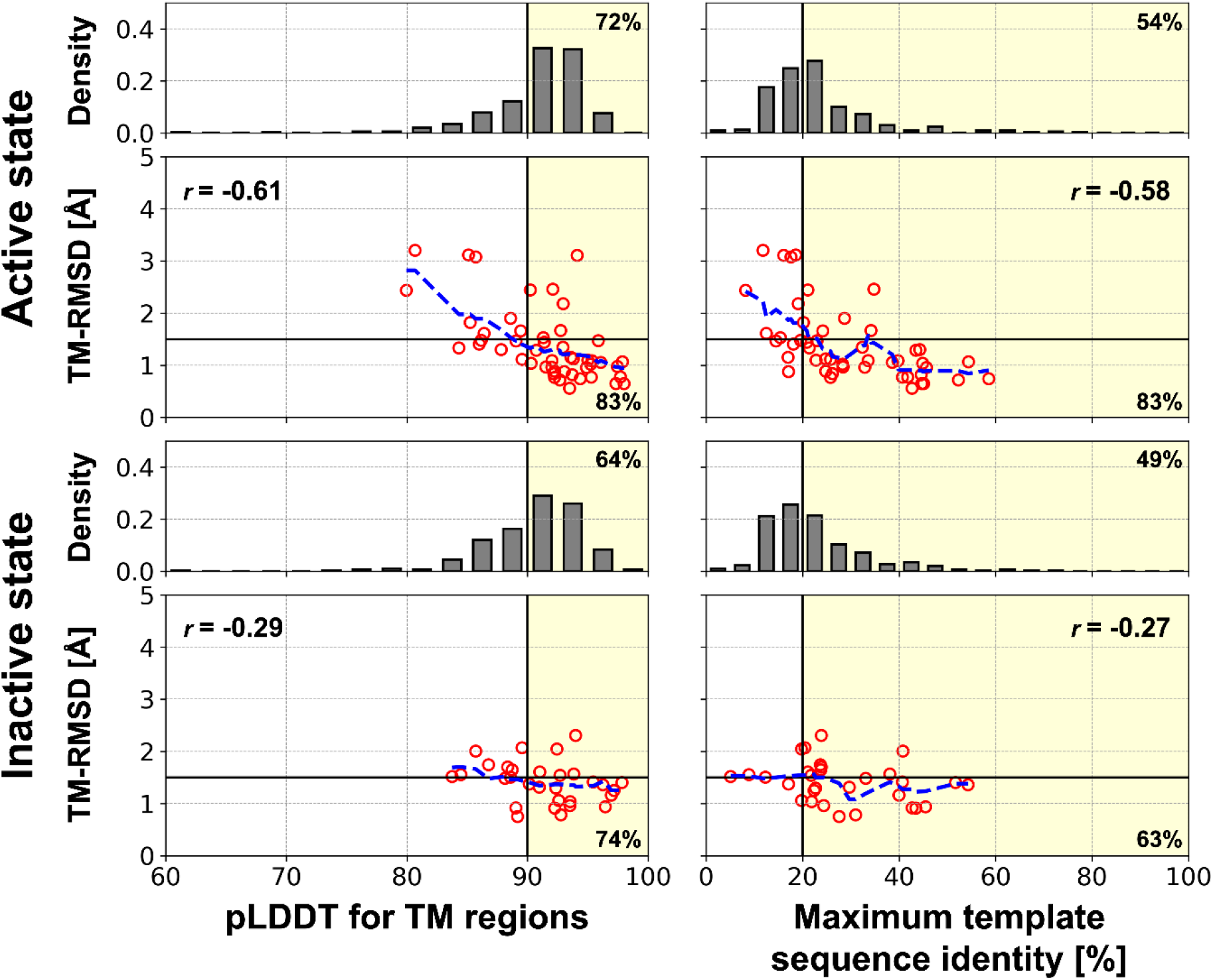
Estimated model accuracies for the multi-state modeling protocol. Relationships between TM-RMSD and the pLDDT or the maximum sequence identity of the structural templates are shown as red circles for the benchmarked GPCR structures. Blue dashed lines represent moving averages of TM-RMSDs with a window of 2.5 for the pLDDT and 5% for the maximum sequence identity. Pearson correlation coefficient for each relationship is shown in the box and denoted as “*r*=“. A model that has a pLDDT higher than 90 or is predicted using a template with a sequence identity of more than 20% was likely to be accurate (TM-RMSD < 1.5 Å). Those selection criteria are shown as yellow background color, and the ratios of accurate models among the selected models are noted at lower right corner. For human GPCRs without experimentally determined structures for each state, distributions of the pLDDT and the maximum sequence identity of the structural templates are shown as grey histograms. The percentages of models that are likely to be accurate assessed by each selection criteria are noted at the top right corner.

### Sampling intermediate conformations

We examined a protocol for sampling intermediate conformations with the human type-1 angiotensin II (AT_1_) receptor (UniProt ID P30556, AGTR1_HUMAN). (**Figure S11**) Rather than utilizing experimental structures in either active or inactive state templates as input structural features, a model generated by linear interpolation between predicted active and inactive state models were fed to the AF2 network model. Input models at different points along the interpolation were used to sample various conformations between the active and inactive states. None of the sequences in the MSA was used except for the target protein sequence. Output models were physically realistic in terms of protein stereochemistry even though they were modeled from unphysical input models. While the input models had continuous structures between the active and inactive models, the output models showed discontinuous structural transitions. (**Figure S11a and b**) Some of these conformations may be considered as representative structures for intermediate states. As we ignored interpolated coordinates for sidechains (except for Cβ atoms), sidechains in intermediate conformations were properly modeled via optimization through the AF2 network. (**Figure S11c**) The sidechain orientations also showed discontinuous transitions. The output models were further validated by mapping the structure onto a potential of mean force (PMF) map generated by a previous MD simulation study.^35^ (**Figure S11d**) The intermediate conformations possessed lower free energy via optimization by the AF2 network from the initially interpolated conformations at higher free energy regions and formed non-trivial pathways. Moreover, some conformations were located at lowest free energy saddle points between the active and intermediate states or between the intermediate and inactive states. It was also possible to generate a few models (colored in cyan in the figure) that closely resemble the intermediate state identified from the MD simulation study.

Recently, Del Alamo *et al.* argued that the AF2 network model can be used for conformational sampling if a shallow MSA is given as input to the model.^36^ Using the protocol, diverse conformations were generated from diverse randomly selected sequences from a deep MSA. It was possible because different combinations of sequences provided different information to the network model and shallow MSAs were not enough to generate a converged structure due to insufficient information. The models also mapped onto higher free energy regions rather than basins, with some models unfolded partially, presumably because of insufficient coevolutionary information from very shallow MSAs with as few as 16 sequences. Randomly selected sequences seem to impose random perturbations to free energy minima structures, which may be analogous to thermal fluctuations around free energy minima structure that can be observed from MD simulations. In contrast, our protocol seems to generate ensemble-averaged models with free energies as low as possible for a given input, *i.e*., different structural features rather than randomness in the MSAs.

We applied both protocols to the multi-state GPCR targets to generate intermediate conformations. (**Figure S12 and S13**) They often generated a range of conformations that spanned between the active and inactive states. However, our protocol occasionally could not generate diverse conformations when active and inactive state models showed little structural difference (*e.g*., CASR/P41180). On the other hand, modeling with shallow MSAs also occasionally failed to sample conformations (*e.g*., MTR1A/P48039) or generated partially unfolded structures (*e.g*., CASR/P41180) when there were very few sequences in MSA. Moreover, in contrast to class A GPCRs, the activation of class B GPCRs involve significant structural changes such as a break of the TM6 helix with a high-energy barrier.^37^ In the case of such complex pathways, it was difficult to capture intermediate states with our protocol (*e.g*., PTH1R/Q03431). In this case, even although input models in our protocol spanned uniformly between the active and inactive state models, the optimized models ended up in either active or inactive state-like conformations with little information about possible intermediate structures.

### Applications of high-accuracy modeling of multiple states

An important question is how higher accuracy translates into practical advantages when models are used further. One application of high-accuracy GPCR models is for the prediction of protein-ligand complexes. GPCRs are one of the major target proteins for approved drugs and remain extremely attractive targets for new drugs.^1,13^ During the initial steps of drug development, computer-aided drug design relies on the prediction of accurate ligand poses and the estimation of ligand binding affinities. This requires the structure of the target protein for being able to dock potential ligands. Because protein-ligand docking is very sensitive to the structure of the binding pocket, high-accuracy models are essential for docking success. In GPCRs the ligand pockets have a different shape depending on its activation state. Therefore, to design a ligand that targets a certain activation state of a GPCR, a high-accuracy protein model for that state is required. To validate the effectiveness of the high-accuracy state-specific GPCR models resulting from the multi-state protocol for protein-ligand docking, ligands from experimental structures were docked to the corresponding GPCR models. (**Figure 4**) Two sidechains at the binding site were set to be flexible because computational model structures were predicted without consideration of binding ligands and resulted in *apo* structures. Otherwise, the success ratios dropped significantly for them because misoriented sidechains prevented a ligand from docking. (**Figure S14**) For active state conformations, using sidechain flexible docking, the protein-ligand docking success with models predicted by our protocol was the highest among all computational models. For around one third of the active state receptors, it was possible to dock ligands within 3 Å in terms of ligand heavy-atom RMSD. However, ligand docking to active state conformations is intrinsically a difficult problem considering that self-docking to less than half of the experimental structures was successful. Docking results for the other computational models were much less satisfactory with less than 10% success. As discussed above, many original AF2 models were more similar to inactive-state structures than active-state structures. and this diminished the accuracy of binding site predictions. (**Figure S15**) Because protein-ligand docking is vulnerable to binding site errors even as low as 1 Å, accurate modeling of the correct activation state structures at the binding site was critical for docking. As an example, in **Figure 4b and c**, an endogenous agonist ligand (serotonin, 5-hydroxytryptamine, SRO from PDB ID 7e2y) is shown after docking to receptor model structures for human 5-hydroxytrypamine receptor 1A (UniProt ID P08908, 5HT1A_HUMAN). The experimental structure of the target protein is in the active state. With a model in the active state from our multi-state modeling protocol, it was possible to dock the ligand with very high accuracy based on a ligand heavy-atom RMSD of 0.55 Å. The protein model has a very accurate binding site structure with 1.05 Å heavy-atom RMSD in the binding site. On the other hand, the AF2 model was modeled in the inactive state, and the docking result to this model was poor with a ligand heavy-atom RMSD of 4.38 Å. As the model was in the inactive state, some of the binding pocket residues were misplaced. Especially W358^6×48^ (indicated by a red arrow in **Figure 4b and c**), which acts as the transmission switch activated by agonist binding, was in its position of inactive state. This misplacement altered the ligand binding pocket, so that the ligand could not be docked into the correct binding pocket.

**Figure 4.**
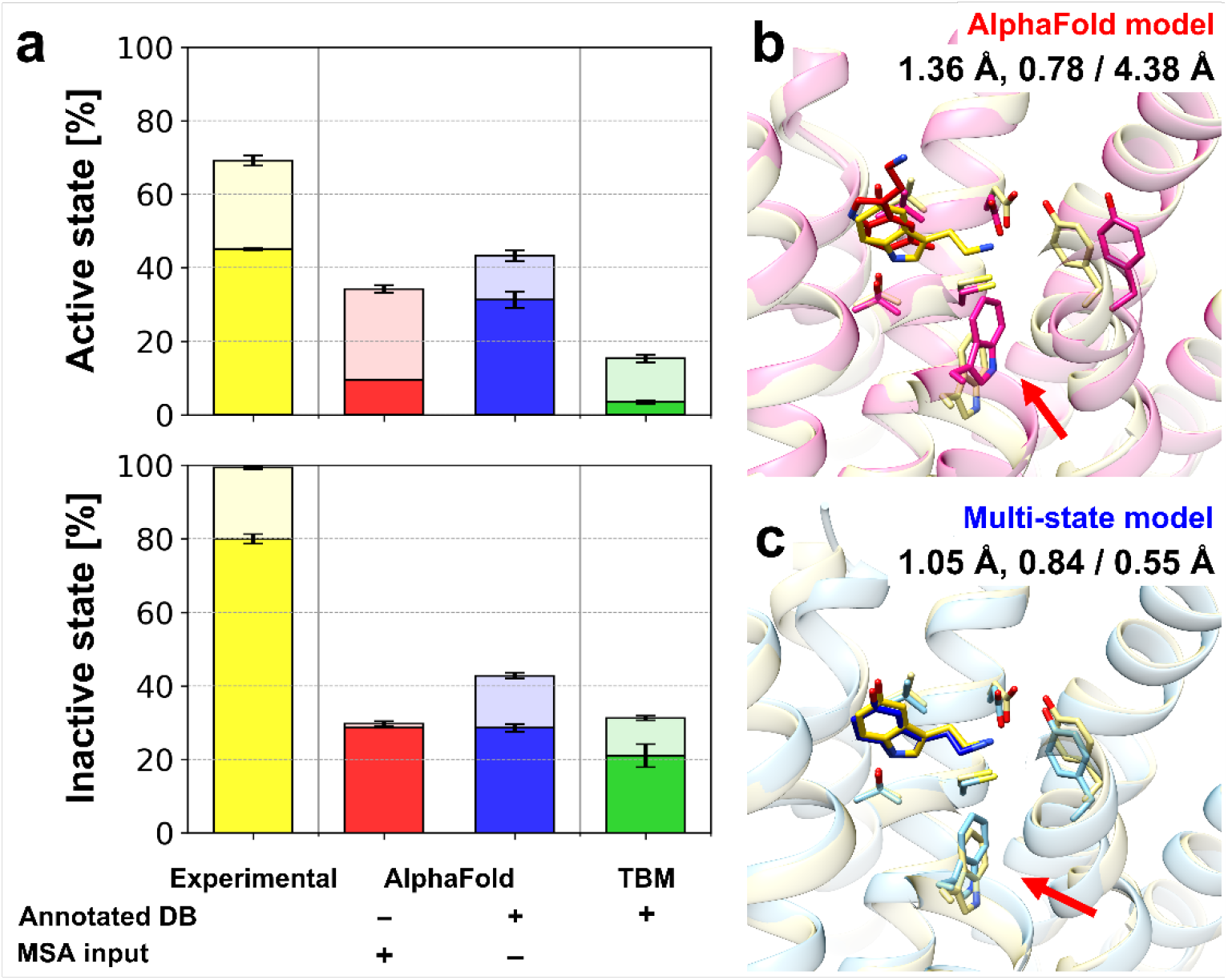
Protein-ligand docking on predicted GPCR models. (a) Protein-ligand docking success ratios for GPCR model structures using various modeling protocols. Docking simulations were performed five times independently for each structure using AutoDock Vina. Success ratios and their standard errors are shown as bar charts for top 1 (opaque) and top 3 (transparent) predictions and overlaid error bars, respectively. An example of protein-ligand docking using different models is presented: (b) the original AlphaFold model and (c) multi-state model. An endogenous agonist ligand (serotonin, 5-hydroxytryptamine, SRO from PDB ID 7E2Y) was docked to protein structure models for human 5-hydroxytrypamine receptor 1A (UniProt ID P08908, 5HT1A_HUMAN). Protein structures are shown in transparent cartoon representations, while docked ligand conformations are depicted as sticks. The experimental protein-ligand complex structure is shown in yellow. Key structural differences between the two protein models that contributed to the different docking performance are indicated by red arrows. The binding site heavy-atom RMSD, the lDDT for the transmembrane region, and the resulting docking accuracy in terms of ligand heavy-atom RMSD are shown under the model names.

For antagonist ligands bound to inactive GPCR states, performance of the protein-ligand docking results with predicted models were similar between our multi-state modeling protocol and the original AF2 with around 30% success, but overall, they were less than 80% success when docking to experimental structures. The lower success rates with inactive state docking for models compared to experimental structures may be because antagonists in the benchmark set are bigger than agonists, so that clashes due to errors at the binding site are more likely.

As another blind test of application of our multi-state modeling, we participated in GPCR Dock 2021. Five GPCR-ligand complexes were given as targets with activation state information. As of March 01, 2022, experimentally determined structures were available for two targets, T02 (GPR139) and T04 (NPY1R), and they were in the active state. For both targets, our protocol predicted more accurate GPCR models than AF2, which predicted models in the inactive state. Especially, for target T04, our active state model was very close to the experimental structure in the active state (PDB ID: 7vgx_R)^38^ with a TM-RMSD of 1.14 Å. (**Figure 5**) Several structural changes upon activation of the protein were successfully modeled. In contrast, AF2 predicted the inactive structure with a TM-RMSD of 0.93 Å with respect to an experimental structure in the inactive state (PDB ID: 5zbh_A), which was included in the AF2 training set, whereas the model deviated from the active state structure with a TM-RMSD of 2.17 Å. Furthermore, our active state model was successful for predicting the complex structure with its endogenous peptide-agonist neuropeptide Y (NPY). (**Figure 5b and c**). The bound peptide was correctly placed with a Cα-RMSD of 2.00 Å. Especially, for the C-terminal residues (residue numbers 32-36), which are important for the binding^38^, our prediction captured the binding pose well with a heavy atom-RMSD of 1.88 Å. As exemplified here, our high accuracy multi-state modeling of GPCRs offers important practical advantages for predicting ligand-bound structures over other methods.

**Figure 5.**
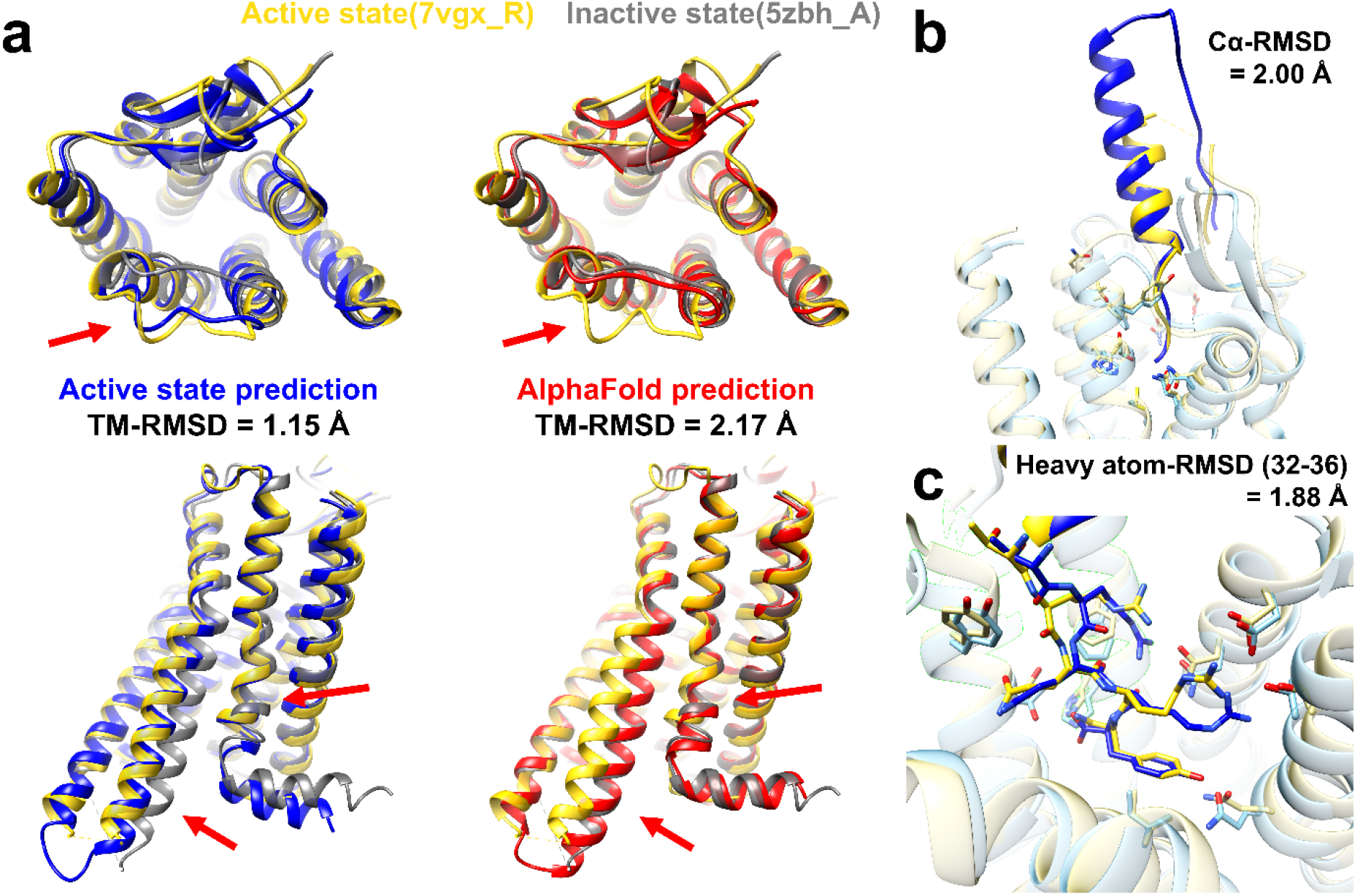
An example of successful GPCR Dock 2021 prediction for target T04, complex of human neuropeptide Y receptor type 1 (UniProt ID P25929, NPY1R_HUMAN) and its endogenous peptide-agonist neuropeptide Y (NPY). The target is in the active state. (a) Active state model using our prediction protocol (blue) and AF2 model (red): (top) view from the extracellular region, (bottom) side-view. Experimental structures of active state (yellow, PDB ID 7vgx_R) and inactive state (gray, PDB ID 5zbh_A) are compared to the predictions. Structural differences due to the activation state difference are pointed by red arrows. (b and c) Neuropeptide Y (NPY) bound to the receptor: (b) overview and (c) focused view on C-terminal residues (residue numbers 32-36). The experimental structure (PDB ID 7vgx_R for the receptor and 7vgx_L for the peptide) and our model are shown in yellow and blue, respectively.

## CONCLUSIONS

Here, we examined AF2 for the modeling of GPCRs in active and inactive conformations. While AF2 clearly generates more accurate models of GPCRs than template-based modeling, the models typically only model one functional state, usually the inactive state, presumably as a result of training based on the PDB where inactive state GPCR structures are overrepresented. A modified protocol is described here that can guide AF2-based high-accuracy modeling towards a specific activation state. In this multi-state modeling protocol, AF2 was used with activation state-annotated GPCR structure databases instead of a general PDB structure database and without MSA input features to avoid learned biases towards inactive GPCR states. With this protocol, it became possible to model both active and inactive states at the same high accuracy as the original AF2 models for just one GPCR state, usually the inactive conformation. The resulting models capture key structural changes upon activation/deactivation of the GPCR at the atomic level. Moreover, we tested a protocol for modeling intermediate states based on interpolated input structures between predicted active and inactive state models. The interpolated structures are used in lieu of templates from a structural database and AF2 network is then used essentially as a refinement program. This resulted in plausible intermediate conformations that may form activation pathways. However, the general applicability of this approach still needs further investigation.

The multi-state models expanded the applicability of AlphaFold. These models improved the correct prediction of ligand binding poses using protein-ligand docking. Especially with active state models, they outperformed other computational models in terms of the docking success rate, reaching almost the same success rate as with experimental structures. Furthermore, GPCR-ligand complex structures were successfully predicted in a blind test, GPCR Dock 2021. The multi-state modeling approach introduced here can be extended in principle to other protein families such as kinases as long as experimental structures in multiple states are available to form state-specific template databases and we expect that the approach described here is a first step towards capturing not just native structures but conformational dynamics at high accuracy via machine learning-based approaches.

## METHODS

### Multi-state protein modeling protocols

Several protein structure prediction protocols were tested to model GPCRs as active and inactive states. First, the original pipeline of AF2^5^ was used to predict GPCR structures. The input consists of target protein sequences, MSAs, and structure templates; as output five models are predicted. We checked whether AF2 can model both active and inactive states. For a GPCR, structural similarities to active and inactive experimental structures were evaluated. The ability of being able to model multiple states was judged based on some of the five models being closer to active state than inactive state while others were closer to the other state. In addition, we intended to guide AF2 to model a specific activation state structure using activation state-annotated GPCR structure databases. A database was built for each active or inactive state. The details about how databases were constructed are described in the following section. Protein structure prediction for a state was guided by the input database for the state, as a replacement of the PDB70 database. ^30^ Databases were used in two ways. One approach was to simply replace the PDB70 database for the structure template search. In another approach, residues in the input MSAs were modified to gaps for sequence positions which were aligned to selected structure templates from the activation state-annotated databases. (**Algorithm S1**) Furthermore, an ablation study was performed to better understand the role of each input feature. The original AF2 method was compared with three variants: one without MSAs, one without structure templates, and one without both MSAs and structure templates. Finally, as a reference, structure prediction was performed via template-based modeling using MODELLER^31^ with the identical structure templates and the sequence alignments used for the AF2 predictions.

### Building activation state-annotated GPCR structure databases

Template structure databases for each active and inactive state GPCRs were built based on the GPCRdb^25,26^ activation state annotation. The database building procedure was based on the official procedure of building customized HHsearch^30^ database. A list of PDB IDs was collected from the GPCRdb for either active or inactive states. GPCR sequences were extracted from mmCIF files of each PDB entry. If there were multiple chains of GPCRs for a PDB entry, a preferred chain selected by the GPCRdb was used. To remove redundant sequences, the extracted sequences were clustered by MMseqs2^39^ with a sequence identity cutoff of 100% and a sequence coverage of 100%. For each representative sequence for a cluster, a multiple sequence alignment was generated using HHblits^30^ by searching homologous sequences against the UniClust30 database with two iterations and HHblits default options. The generated multiple sequence alignments were post-processed to build a HHsearch database for a given state. As of July 29, 2021, there were 224 and 309 experimentally determined structures for active and inactive state structures, respectively. And they resulted in 161 and 206 unique entries for the activation state-annotated GPCR structure databases after removing proteins with identical sequences.

### Benchmark tests

A benchmark set of human GPCRs was used to evaluate the different protocols for multi-state modeling of proteins. The set was composed of human GPCRs that were experimentally first determined after May 01, 2018 and before January 05, 2022. Since AF2^5^ was trained protein structures determined by April 30, 2018, thus, none of the GPCR structures that were used for the training were included in the benchmark set. There were 68 GPCRs in the set, and they are summarized in **Table S2**. Among them, there were 49 and 30 GPCRs for the active and inactive states, and 15 GPCRs were determined in both active and inactive states. For the benchmark test, close homologous structures that have a sequence identity higher than a cutoff of 70% were excluded from the template lists. For the model accuracy evaluation in terms of TM-RMSD, the Cα-RMSD was evaluated using transmembrane helices, whose definition was taken from the GPCRdb.^25,26^ An experimental structure was considered as a reference structure if it had a resolution of 4.0 Å or higher. If there were multiple experimental structures for an activation state of a GPCR, the closest result was reported. For the evaluation of loop model qualities, Cα-RMSDs of three intracellular loops (ICLs) and three extracellular loops (ECLs) were calculated. A loop is defined as a region from the last residue of a TM helix to the first residue of its next TM helix. Loop residues that have B-factors higher than 100 Å^2^ were excluded from the loop analysis because of the associated structural uncertainties.

All human non-olfactory GPCRs were modeled as active and inactive state structures. The list of the GPCRs was retrieved from GPCRdb.^25,26^ (https://gpcrdb.org/alignment/targetselection) Since some of the GPCRs have non-transmembrane domains, we modeled the transmembrane (TM) domain only to focus on its activation state. The UniProt topology annotations were used to define the TM domains. Up to 100 residues towards N- and C-termini from the first and seventh TM helices were additionally considered for modeling not to lose any of their interactions with the TM domain and relevant TM helix residues because the annotations are based on a TMHMM prediction.^40^

### Sampling intermediate conformations

Intermediate conformations are modeled based on the identical approach that we used for modeling of the active and inactive state models. Rather than searching structural templates against activation state-annotated GPCR databases, artificial input templates were generated and were fed to the AF2 network model. After superposing predicted active and inactive state models, atom coordinates were linearly interpolated between the models with ranges of the degree of activation, *d* (**Eq. 1**).

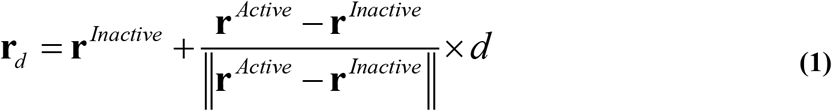

In this work, we interpolated backbone atoms (N, Cα, C, and O) and Cβ atoms (for non-Glycine residues), so that sidechains were freely optimized by the AF2 network model according to changes in their neighboring atoms. For the degree of activation, *d*, we used equally spaced values from 0 to 100 % with an incremental of 5 %; it resulted in 21 input models. For each input model with a different degree of activation, one output model was predicted by the AF2 network model.

Various conformations were sampled by following the protocol described in Del Alamo et al.^36^ For a target GPCR, a MSA was generated by the ColabFold MMseqs2 API.^41^ From the original MSA, shallow MSAs were subsampled by modifying the AF2 network parameters, max_msa_clusters and max_extra_msa. In this work, according to the suggested values in the original work, we used 16, 32, 64, and 128 for max_extra_msa and set max_msa_cluster to half of the value. For each depth of MSA, twenty models were generated for a target with different random seeds.

### Protein-ligand docking using AutoDock Vina

To validate the usefulness of high-accuracy GPCR models, protein-ligand docking tests were performed. Target GPCR-ligand complexes were among the experimental structures for the benchmark set described above. We selected ligands that have not too many rotatable torsion angles (< 15) and heavy atoms (< 25) because the docking procedure described below may not effectively handle such high degrees of freedom and lead to docking failures. The tested GPCR-ligand complexes are summarized in **Table S3**. As protein structures, we used the original AF2 models, multi-state models in the corresponding activation states that of the experimental structures, and template-based models in addition to the experimental structures. Protein-ligand docking was performed five times independently for a given protein structure. Ligand heavy-atom RMSDs with consideration of symmetry were calculated to evaluate the accuracy of protein-ligand docking using GalaxyDock.^42^ A docking simulation was considered successful if the ligand heavy-atom RMSD was lower than 3 Å.

AutoDock Vina^43^ was used to perform protein-ligand docking on the predicted GPCR structures. Protein structures were prepared using the “prepare_receptor’ tool of the ADFR suite.^44^ Ligand structures were extracted from the experimental structures and prepared for docking. They were first converted to the Tripos Mol2 File format by adding hydrogens at pH=7 and converted to the PDBQT file for AutoDock Vina. All torsion angles were set to be rotatable except for torsion angles around amide and guanidinium bonds, which are the default option for the “prepare_ligand” tool of ADFR suite. In addition to the ligand flexibility, two protein sidechains were set to be flexible to consider conformational changes upon ligand binding. These sidechains were selected among the binding pocket sidechains that were in close contact with the ligand (heavy-atom distance < 3 Å). If there were more than two sidechains met the criteria, the two closest ones were selected. Atomic charges of ligand atoms were assigned using the Gasteiger charge model.^45^ Protein-ligand docking was carried out at the binding pocket for a ligand. For an experimental protein structure, the center of the cubic search space of docking was located at the geometrical center of the ligand from its bound structure, and the width of the search space was 20 Å for each axis. For predicted GPCR structures, the same search space was used after superimposition to the experimental structure. Protein-ligand docking was performed using AutoDock Vina with an exhaustiveness of 32 and repeated five times for a protein structure and a ligand pair.

The protein-ligand docking success ratio was related to protein model qualities for each protein structure prediction protocol and each activation state of the GPCR. We used TM-RMSD and lDDT^46^ as the overall protein model accuracy metrics for backbone atoms and additional consideration of sidechains, respectively. In addition to them, the binding site heavy atom-RMSD was used to measure the binding site accuracy. The binding site residues were defined as residues whose atoms were within 8 Å from the ligand in the experimental structure.

### GPCRDock2021 using multi-state prediction protocol

GPCR-ligand complex structures for GPCRDock2021 were predicted using a two-stage modeling protocol. First, receptor structures were modeled using the multi-state prediction protocol. The activation state of target GPCRs were determined based on the given information of G-protein binding. Then, GPCR-ligand complexes were modeled using AutoDock Vina for organic ligand molecules or AlphaFold-Multimer^47^ for peptide ligands. For organic ligand molecules, we mostly followed the docking protocol described above except for ligand structure preparation and positioning of the docking search space. Ligand structures were generated from SMILES strings using UCSF Chimera.^48^ The center of the docking search space was manually set at the canonical class A GPCR binding pocket. For peptide ligands, we utilized a modified AlphaFold-Multimer protocol. The modification enabled AlphaFold-Multimer model to take any structure as a template rather than performing template search so that we could feed the predicted GPCR structures as input. We did not use our multi-state prediction protocol for peptide docking because that approach removes MSA information while coevolutionary information from MSA was essential for predicting inter-protein contacts.

## Availability

The activation annotated GPCR databases and predicted models in either activation states are deposited at https://doi.org/10.5281/zenodo.5745217. The scripts used for the modeling protocol are shared at https://github.com/huhlim/alphafold-multistate. A simplified protocol using ColabFold^41^ is also available at https://colab.research.google.com/github/huhlim/alphafold-multistate/blob/main/AlphaFold_multistate.ipynb.

## ACKNOWLEDGEMENT

This research was supported by National Institutes of Health Grant R35 GM126948. We appreciate Martin Steinegger for building and utilizing customized HHsearch databases and Gyu Rie Lee for helpful discussions for modeling GPCRs.

## AUTHOR CONTRIBUTIONS

LH and MF designed the research, LH performed and analyzed the work, and LH and MF jointly wrote the manuscript.

