## supplementary figures and tables for "Multi-State Modeling of G-protein Coupled Receptors at Experimental Accuracy"

*Corresponding author:

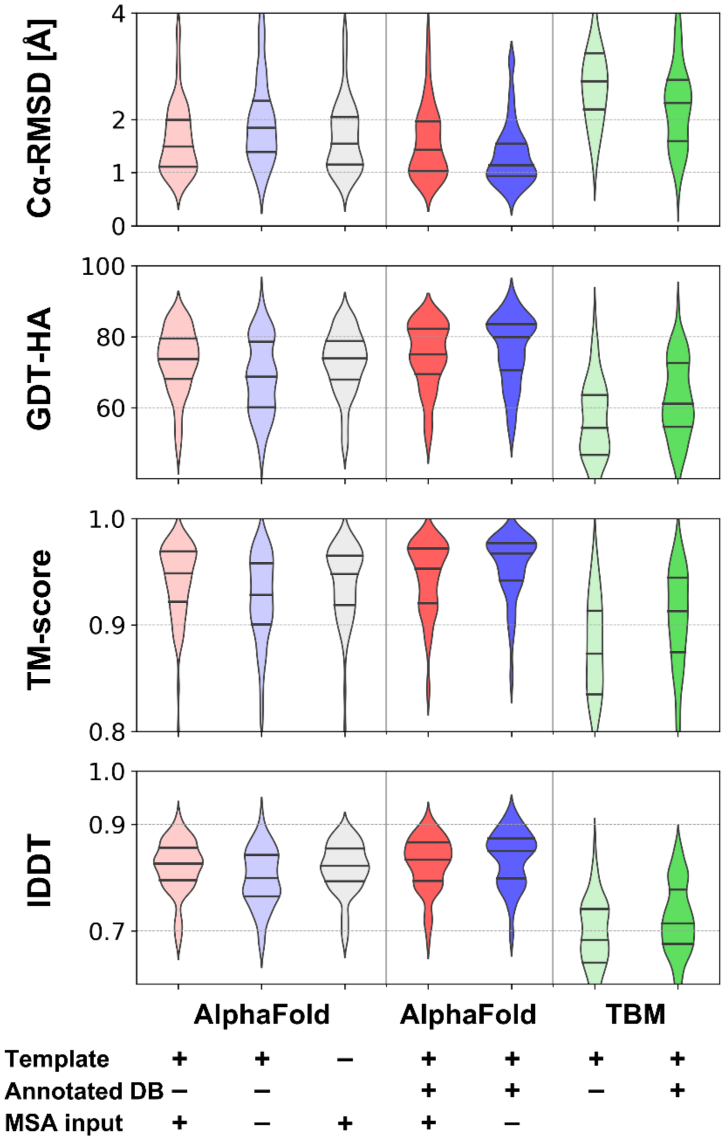


**Figure S1.** Modeling accuracy for GPCRs using various modeling protocols based on AlphaFold and template-based modeling (TBM) regardless of their activation state. Accuracies are measured on transmembrane helix regions only. If there are multiple experimental structures for a GPCR, the best structural accuracy was used for an evaluation of a model. Distributions of modeling accuracy are shown as violin plots, and three quartiles are shown as black lines.

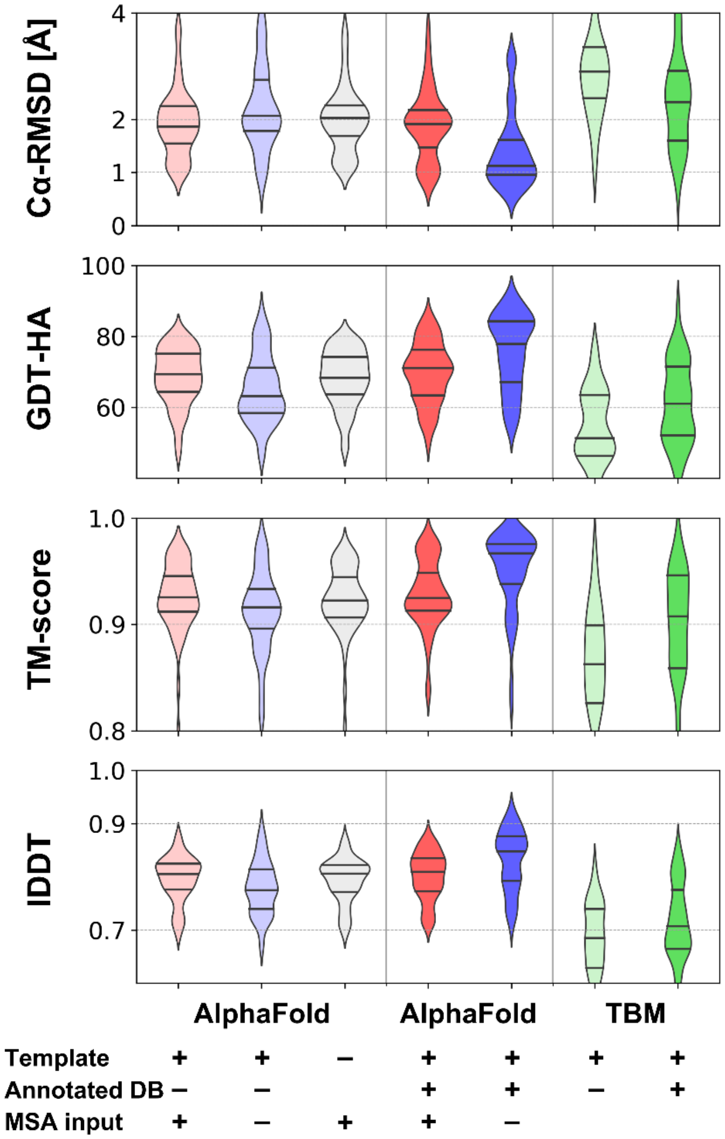


**Figure S2.** Modeling accuracy for the active state GPCRs using various modeling protocols. See **Figure S1** for details.

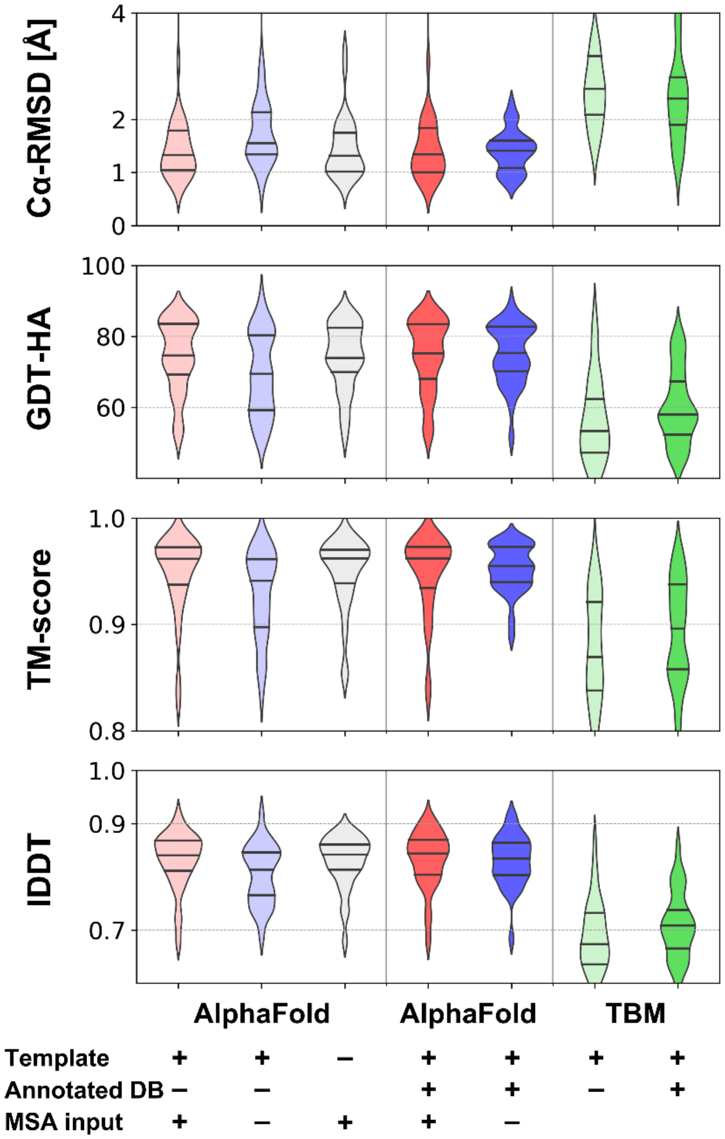


**Figure S3.** Modeling accuracy for the inactive state GPCRs using various modeling protocols. See **Figure S1** for details.

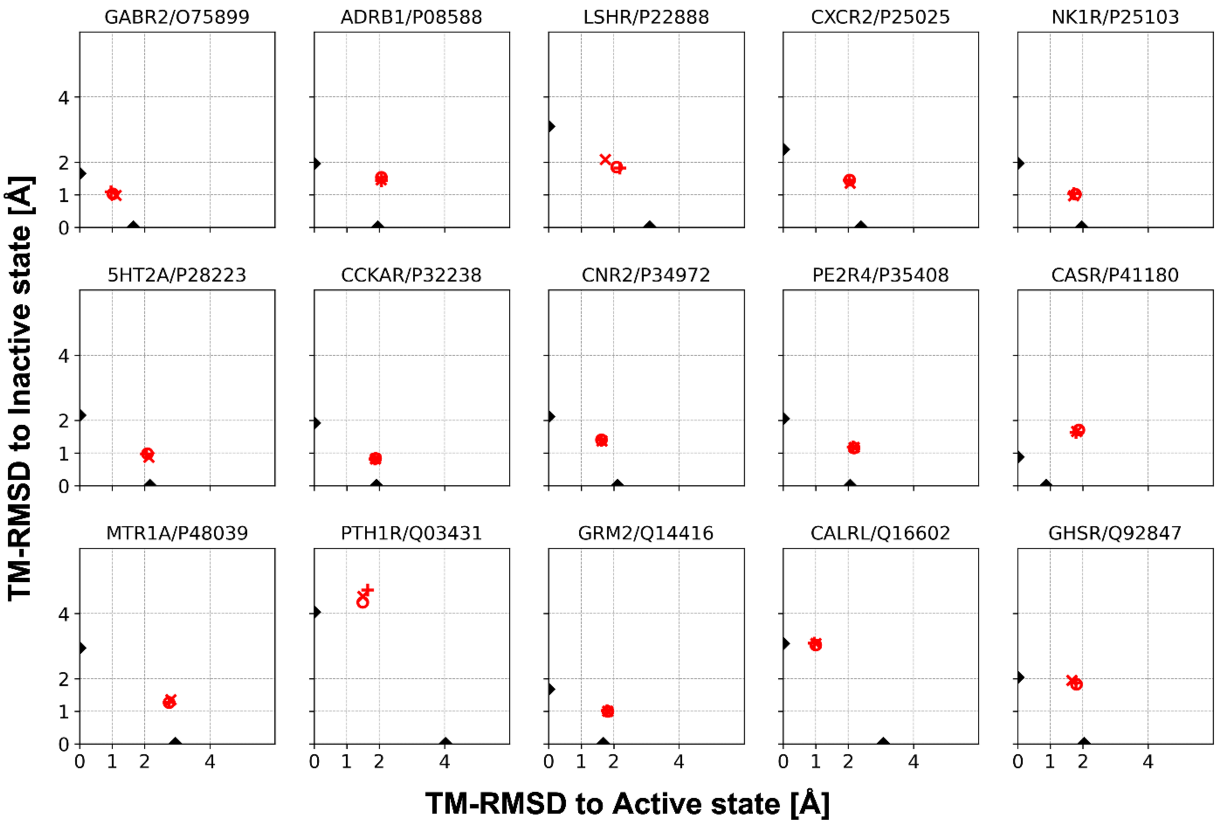


**Figure S4.** Structure similarities of AlphaFold2 models with MSA input features. TM-RMSDs are measured with respect to active and inactive states of individual human GPCRs. Models predicted with the standard PDB70 database are shown as plus signs. Models predicted with state-annotated GPCR databases for active and inactive states are shown as circles and Xs, respectively. Structure similarities between active and inactive state experimental structures are shown as black triangles on both axes.

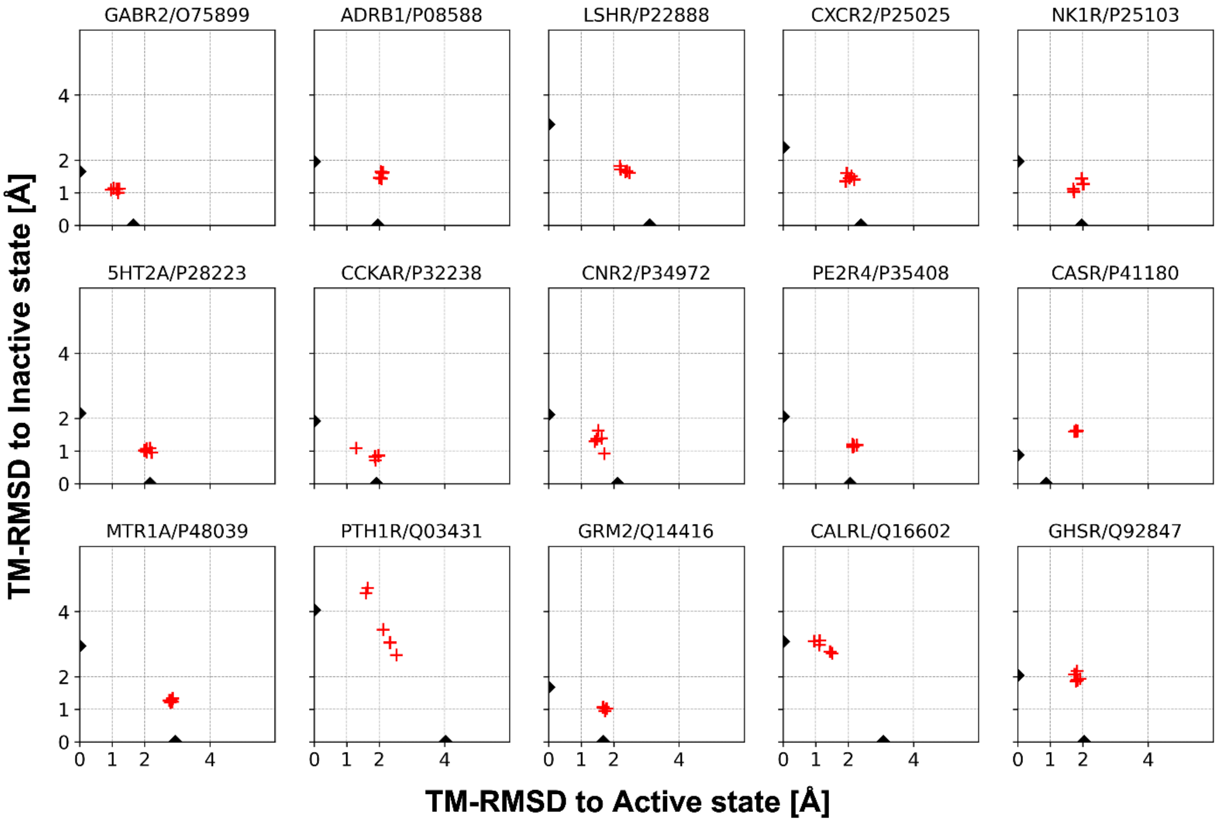


**Figure S5**. Structure similarities of five models of the original AlphaFold2. TM-RMSDs are measured with respect to active and inactive states of individual human GPCRs. Structure similarities between active and inactive state experimental structures are shown as black triangles on both axes.

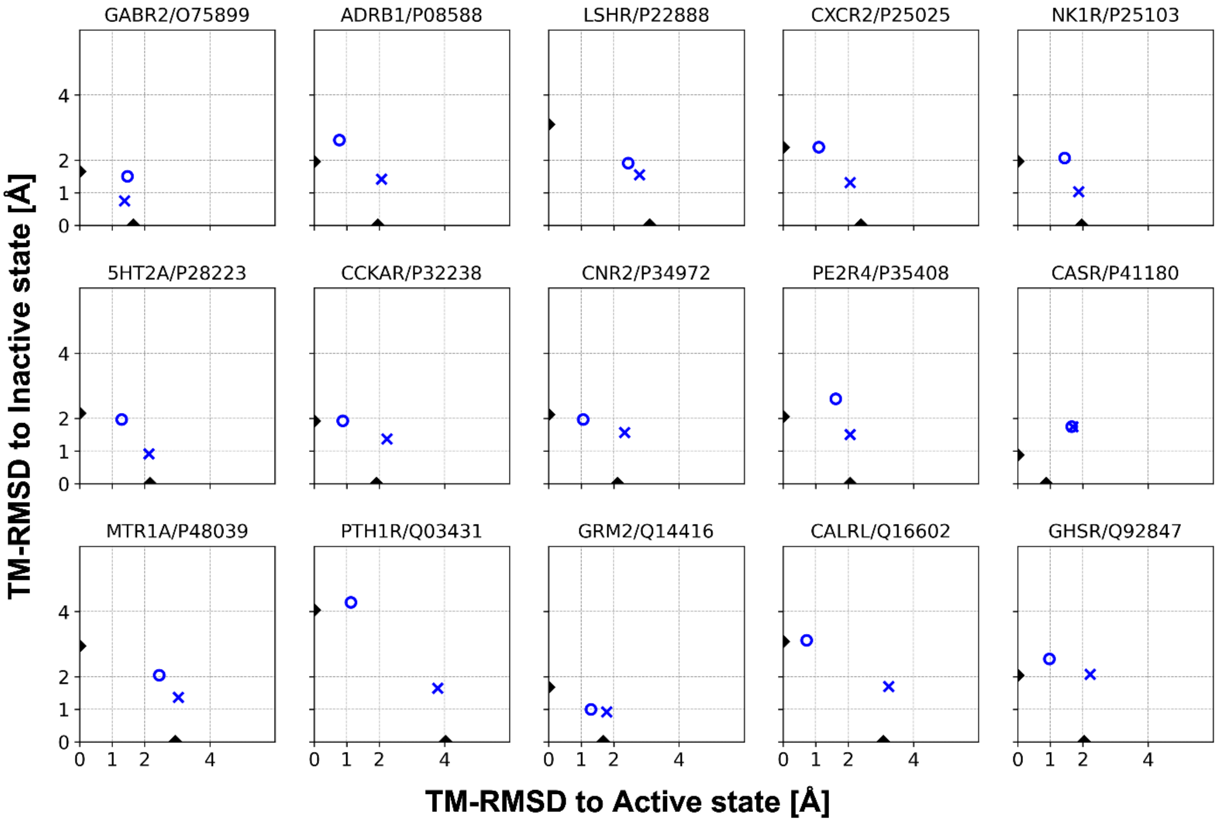


**Figure S6.** Structure similarities of AlphaFold2 models without MSA input features. See **Figure S4** for details.

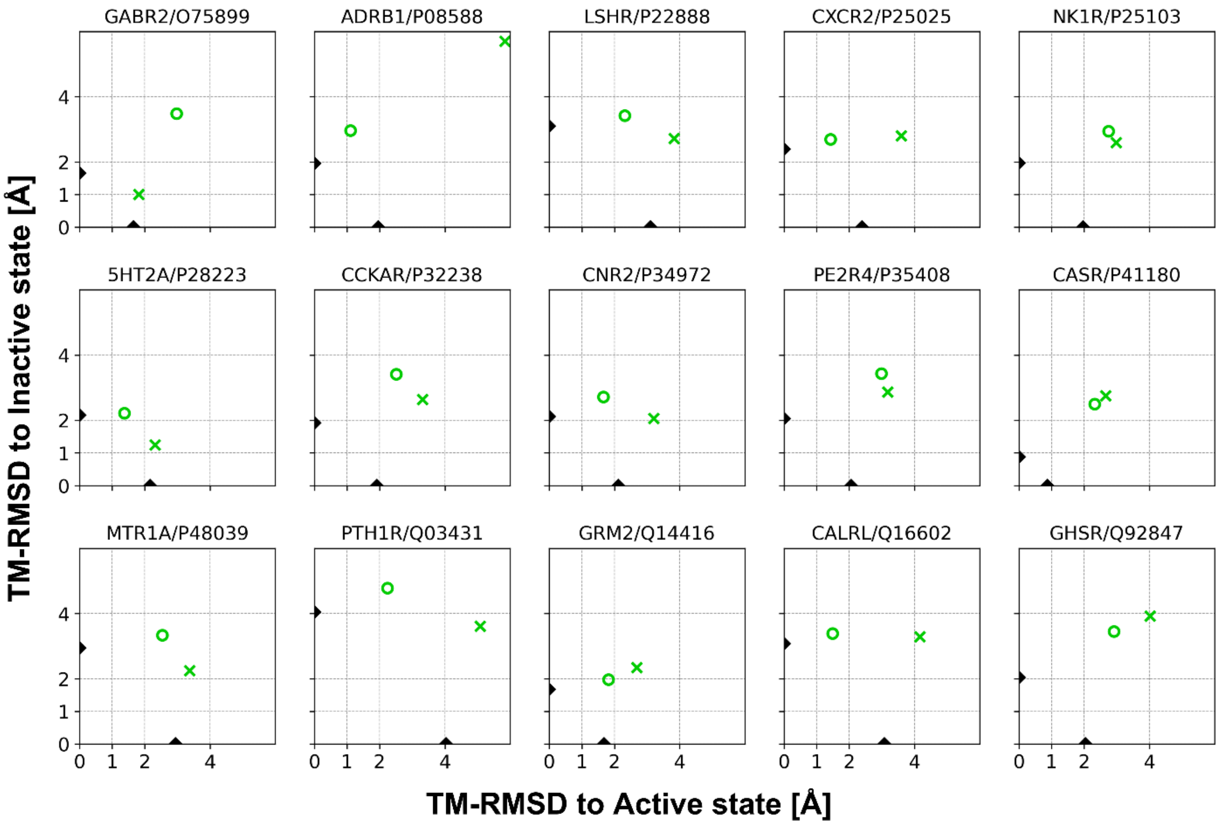


**Figure S7.** Structure similarities of template-based models. See **Figure S4** for details.

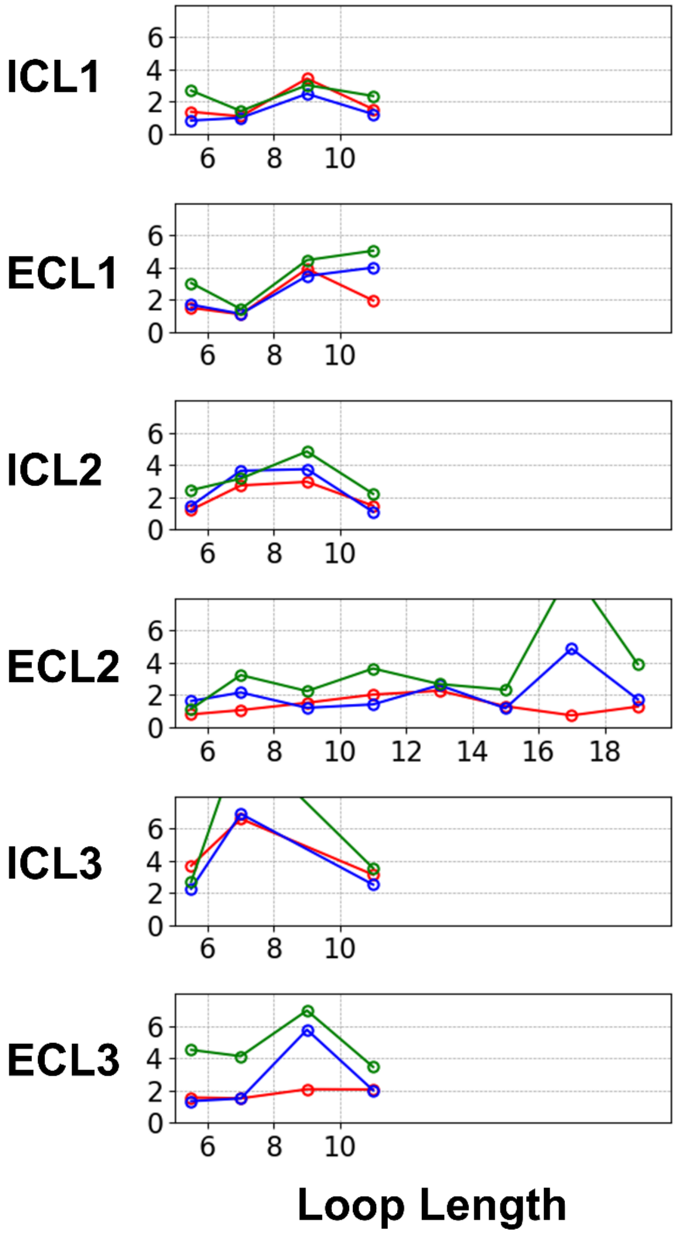


**Figure S8.** Loop model qualities in Cα-RMSD [Å] as a function of the loop length. Model accuracies for the original AlphaFold2, our multi-state modeling protocol, and template-based modeling are shown in red, blue, and green, respectively.

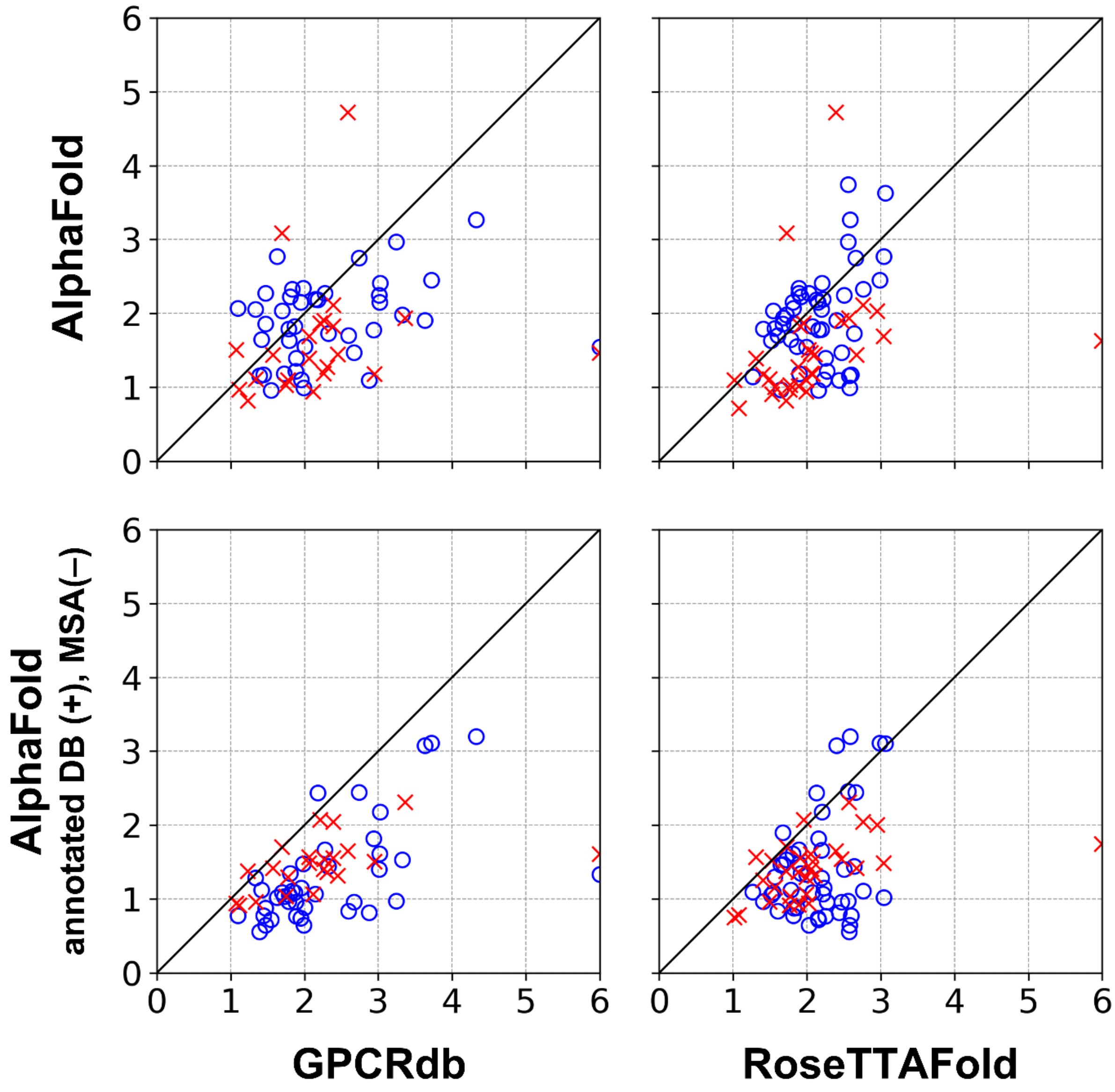


**Figure S9.** Modeling accuracy comparisons with other modeling methods that are able to predict in a specific activation state in terms of TM-RMSD [Å]. The GPCRdb models were retrieved from a GPCRdb archive (2018 April). RoseTTAFold models were generated using the same structural templates that were used for our multi-state AlphaFold modeling protocol. Data for the active and inactive states are represented in blue circles and red Xs, respectively. Outlier data points are clipped at the edge of it.

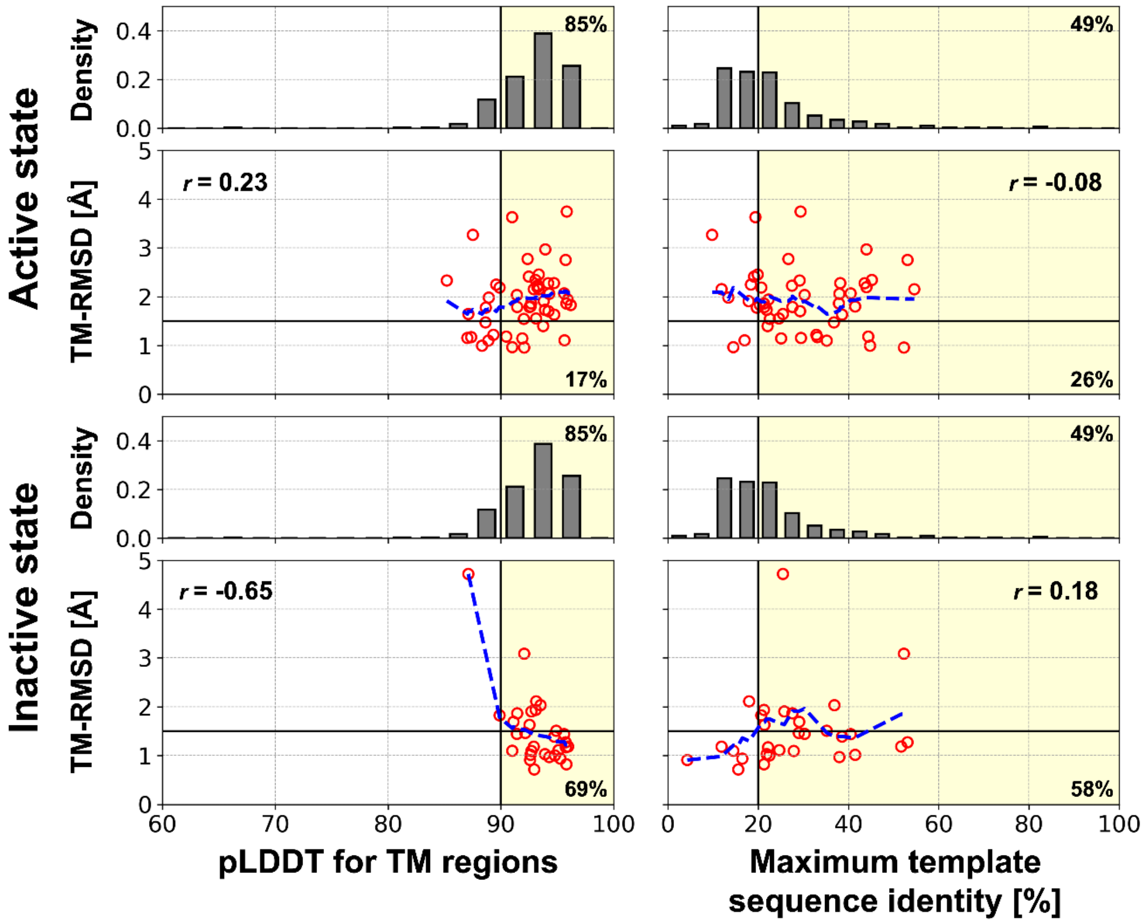


**Figure S10.** Estimation of model accuracies for the original AF2 protocol. See **Figure 3** for details.

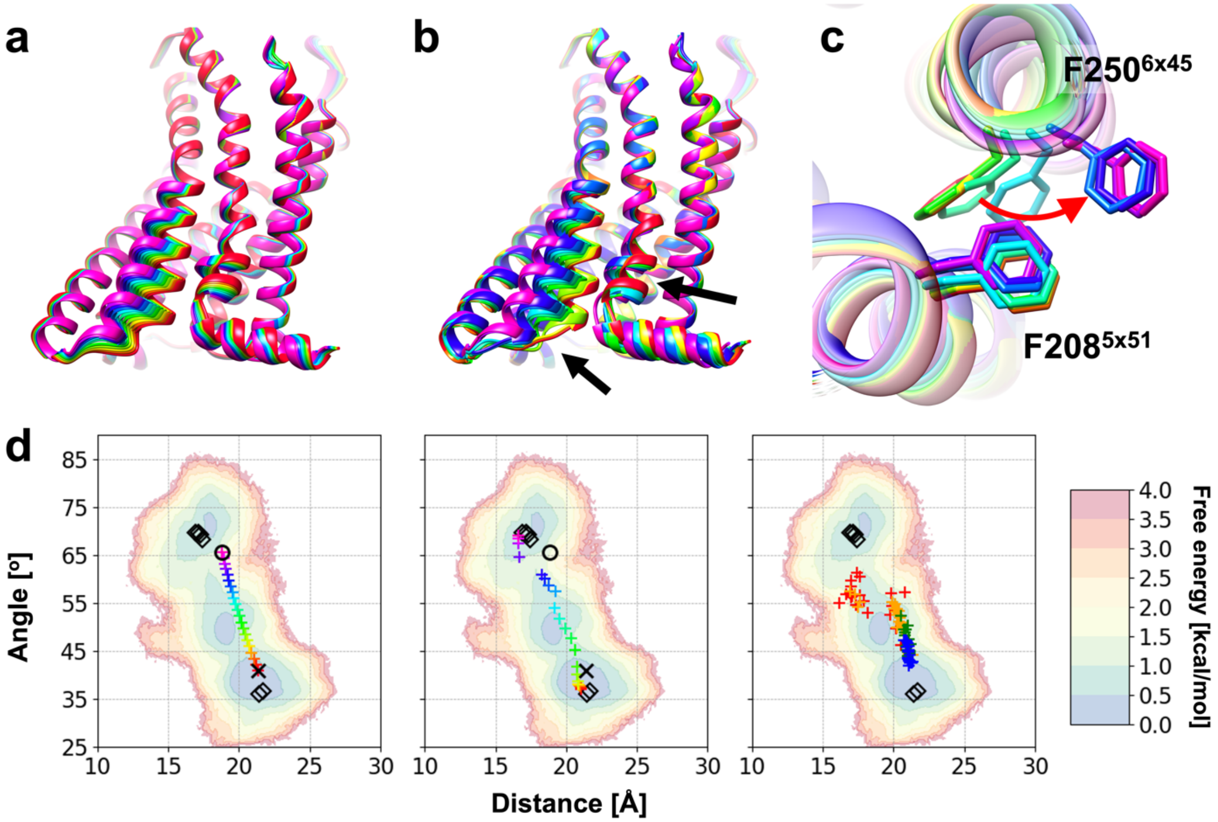


**Figure S11.** Sampling of intermediate conformations for the human type-1 angiotensin II ($AT_1$) receptor (UniProt ID P30556, AGTR1_HUMAN) (a) Input models, which were generated by interpolating between the active and inactive state models. They are depicted in ranges of colors from red (inactive) to purple (active). (b) Output models, which are presented in the same color of the corresponding input models. For brevity, every two models are presented among twenty-one input/output models. Representative structural changes upon activation are pointed out by black arrows. (c) Atomistic details of one key structural transition, $F^{6x45}$ ratchet past $F^{5x51}$. The direction of the transition upon activation is shown with a red arrow. (d) Sampled models mapped onto a potential of mean force (PMF) map generated by a molecular dynamics simulation study.[1] Conformational landscape of the receptor is described by a Cα atom distance between $L^{5x55}$ and $N^{7x46}$ and an angle among $F^{6x34}$, $S^{6x47}$, and $V^{2x41}$ Cα atoms. The PMF is depicted as a contour plot, and the corresponding color scales are shown as a color bar on the right. Experimentally determined structures are mapped as black diamonds. (left and middle) Active and inactive state models by our multi-state modeling protocol are shown as circles and Xs, respectively. Input models (left) and output models (middle) are shown as plus signs with the same color scheme for panels a and b. (right) Models generated using shallow MSAs: 16 (red), 32 (orange), 64 (green), and 128 (blue) sequences.[2]

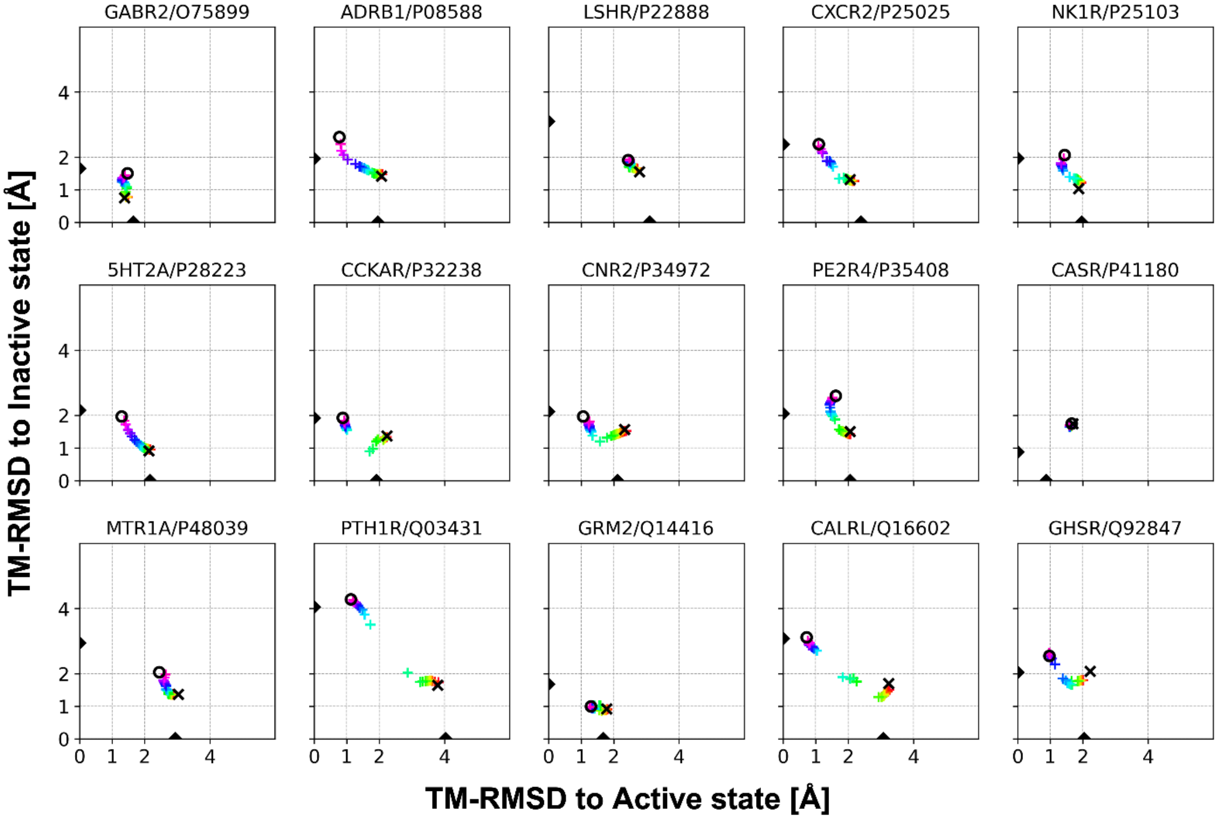


**Figure S12.** Structure similarities of AlphaFold2 models with sampled intermediate conformations using interpolated input structures. Two end points, the active and inactive state models predicted by our multi-state modeling protocol, are shown as black circles and Xs, respectively. Twenty-one output models are shown as plus signs, and the colors represents the degree of activation of the input interpolated structures with ranges of colors from red (inactive) to purple (active). Structure similarities between active and inactive state experimental structures are shown as black triangles on both axes.

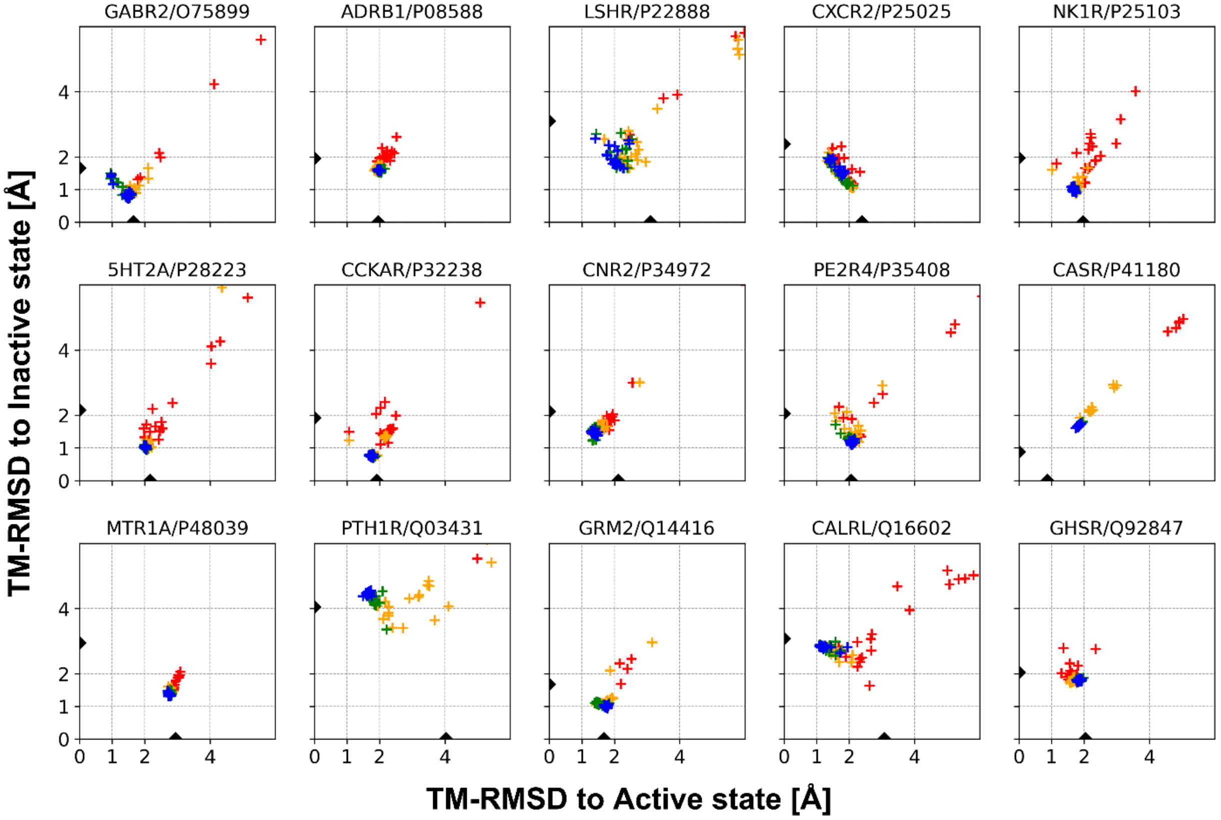


**Figure S13.** Structure similarities of AlphaFold2 models with shallow MSAs. The maximum number of sequences in the MSAs were 16 (red), 32 (orange), 64 (green), and 128 (blue). Twenty models for a maximum number of sequences for each target were generated with different random seeds. Structure similarities between active and inactive state experimental structures are shown as black triangles on both axes.

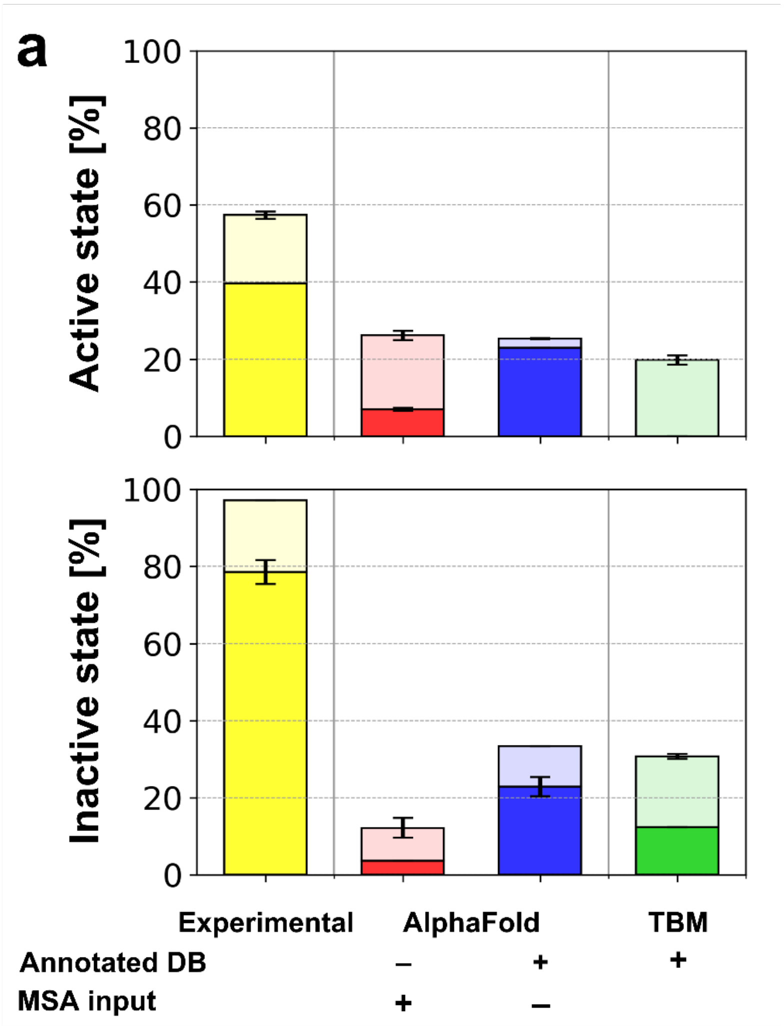


**Figure S14.** Protein-ligand rigid-body docking success ratios for GPCR model structures using various modeling protocols. See **Figure 4** for details.

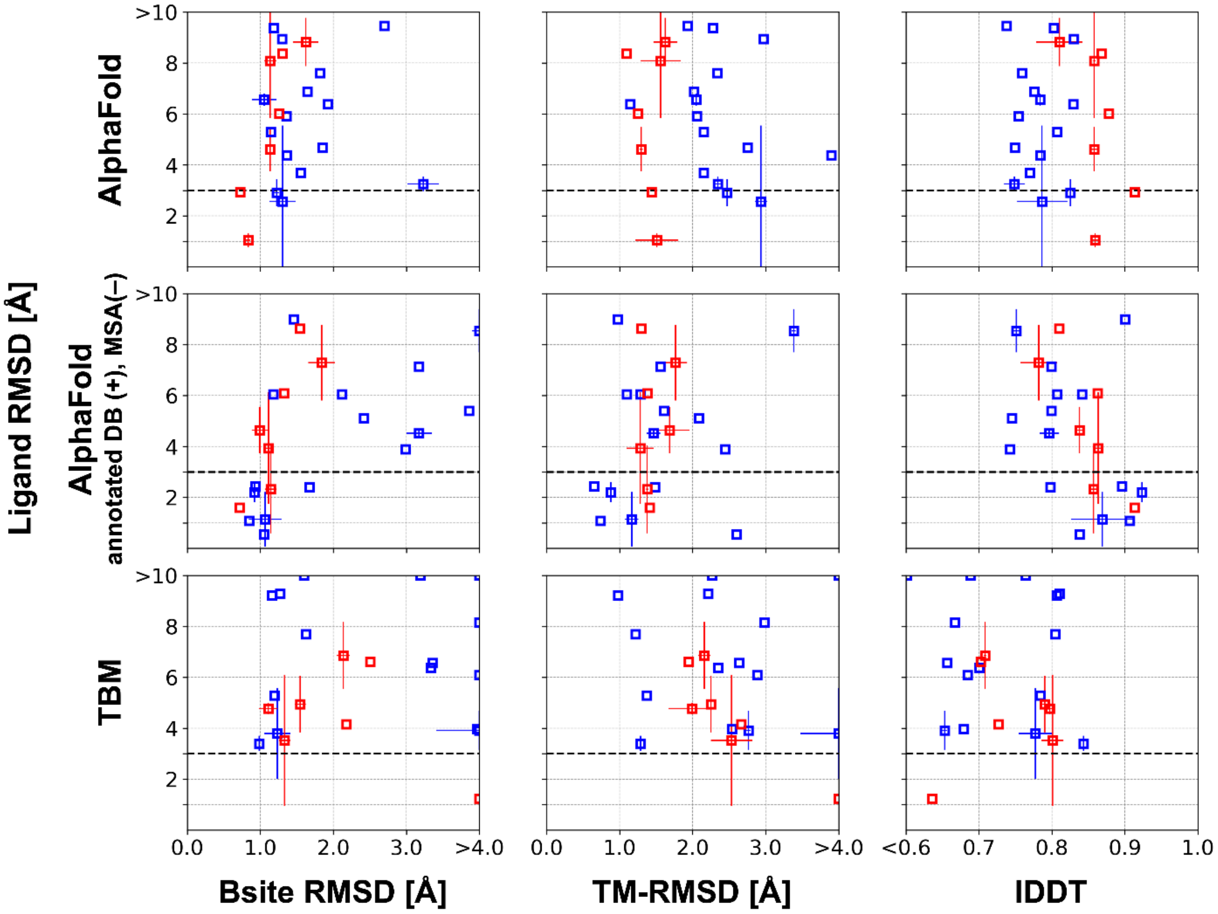


**Figure S15.** Relationships between the receptor structure accuracy and the protein-ligand docking accuracy. Data for the active and the inactive state protein-ligand complexes are shown in blue and red, respectively. Each data point represents the average quality of protein-ligand docking results for multiple ligands that docked to a GPCR model. Outlier data points are clipped at the edge of it.

**Table S1.** GPCR structure modeling accuracy in TM-RMSD [Å] by GPCR classes.[1]

| **Overall** | **All** | **A** | **B1** | **B2** | **C** | **F** |
|---|---|---|---|---|---|---|
| AF2[2] | 1.490<br>(34/68, 50%) | 1.573<br>(20/47, 43%) | 1.176<br>(8/10, 80%) | 3.266<br>(0/1, 0%) | **1.014**<br>(4/7, 57%) | **1.168**<br>(2/3, 67%) |
| Multi-state modeling[3] | **1.137**<br>(47/68, 69%) | **1.332**<br>(31/47, 66%) | **0.888**<br>(10/10, 100%) | **2.608**<br>(0/1, 0%) | 1.516<br>(3/7, 43%) | 1.250<br>(3/3, 100%) |
| TBM[3] | 2.306<br>(14/68, 21%) | 2.511<br>(8/47, 17%) | 1.557<br>(4/10, 40%) | 3.529<br>(0/1, 0%) | 2.084<br>(2/7, 29%) | 1.880<br>(0/3, 0%) |
| **Active state** | **All** | **A** | **B1** | **B2** | **C** | **F** |
| AF2[2] | 1.862<br>(12/49, 24%) | 2.067<br>(3/33, 9%) | 1.176<br>(8/10, 80%) | 3.266<br>(0/1, 0%) | 1.793<br>(1/4, 25%) | 1.941<br>(0/1, 0%) |
| Multi-state modeling[3] | **1.122**<br>(35/49, 71%) | **1.287**<br>(22/33, 67%) | **0.888**<br>(10/10, 100%) | **3.202**<br>(0/1, 0%) | **1.562**<br>(2/4, 50%) | **1.471**<br>(1/1, 100%) |
| TBM[3] | 2.324<br>(10/49, 20%) | 2.615<br>(6/33, 18%) | 1.557<br>(4/10, 40%) | 3.941<br>(0/1, 0%) | 2.647<br>(0/4, 0%) | 2.303<br>(0/1, 0%) |
| **Inactive state** | **All** | **A** | **B1** | **B2** | **C** | **F** |
| AF2[2] | **1.329**<br>(19/30, 63%) | 1.413<br>(13/20, 65%) | 3.902<br>(0/2, 0%) | - | **1.055**<br>(4/6, 67%) | **1.084**<br>(2/2, 100%) |
| Multi-state modeling[3] | 1.407<br>(17/30, 57%) | **1.407**<br>(12/20, 60%) | **1.672**<br>(0/2, 0%) | - | 1.216<br>(3/6, 50%) | 1.205<br>(2/2, 100%) |
| TBM[3] | 2.389<br>(5/30, 17%) | 2.529<br>(3/20, 15%) | 3.444<br>(0/2, 0%) | - | 2.211<br>(2/6, 33%) | 1.712<br>(0/2, 0%) |

[1] The median TM-RMSD for each GPCR class and each modeling protocol are presented and highlighted in red if it is better than 1.5 Å. Best performance among three protocols were in bold characters. The number of high-accuracy predictions (TM-RMSD < 1.5 Å), total number of GPCRs for each class, and the percentage of high-accuracy predictions are shown in the parentheses.

[2] The standard PDB70 database was used for the template database.

[3] Activation state-annotated GPCR databases were used for the template database.

**Table S2.** Information of the recently experimentally determined human GPCR set.

| UniProt ID[1] | Gene name[1] | Class | Number of PDB structures[2] | Number of MSA sequences | Maximum template sequence identity [%][3] |
|---|---|---|---|---|---|
| P08908 | 5HT1A | A | 3/3/0 | 142976 | 29/34/37 |
| P28221 | 5HT1D | A | 1/1/0 | 129145 | 54/58/54 |
| P28566 | 5HT1E | A | 1/1/0 | 107056 | 43/44/45 |
| P30939 | 5HT1F | A | 1/1/0 | 115134 | 44/54/44 |
| **P28223** | 5HT2A | A | 7/1/6 | 116856 | 38/43/43 |
| P08912 | ACM5 | A | 1/0/1 | 111796 | 35/35/45 |
| P08913 | ADA2A | A | 2/0/2 | 127200 | 29/43/21 |
| P18089 | ADA2B | A | 1/1/0 | 133186 | 44/28/46 |
| P18825 | ADA2C | A | 1/0/1 | 119106 | 27/28/22 |
| **P08588** | ADRB1 | A | 4/3/1 | 123468 | 40/40/40 |
| P46663 | BKRB1 | A | 1/1/0 | 74174 | 22/25/22 |
| P30411 | BKRB2 | A | 1/1/0 | 78880 | 24/24/22 |
| **P32238** | CCKAR | A | 8/5/3 | 138602 | 21/17/17 |
| P51684 | CCR6 | A | 1/1/0 | 72743 | 27/32/29 |
| P32248 | CCR7 | A | 1/0/1 | 66412 | 29/34/33 |
| Q9Y271 | CLTR1 | A | 2/0/0 | 64264 | 28/23/25 |
| Q9NS75 | CLTR2 | A | 4/0/0 | 64091 | 25/22/26 |
| **P34972** | CNR2 | A | 4/2/2 | 30085 | 38/38/38 |
| **P25025** | CXCR2 | A | 3/2/1 | 85793 | 30/33/29 |
| P21728 | DRD1 | A | 11/11/0 | 93744 | 26/28/26 |
| P25090 | FPR2 | A | 2/2/0 | 63977 | 29/26/29 |
| P32239 | GASR | A | 2/2/0 | 129190 | 20/39/16 |
| **Q92847** | GHSR | A | 5/4/1 | 99833 | 27/28/20 |
| P30968 | GNRHR | A | 1/0/1 | 30850 | 18/19/19 |
| Q6DWJ6 | GP139 | A | 4/4/0 | 11094 | 18/18/15 |
| Q8TDU6 | GPBAR | A | 3/3/0 | 2112 | 19/16/16 |
| Q9Y2T5 | GPR52 | A | 4/1/0 | 38031 | 19/20/19 |
| **P22888** | LSHR | A | 3/2/1 | 22406 | 20/8/8 |
| Q15722 | LT4R1 | A | 1/0/1 | 69829 | 21/23/23 |
| P32245 | MC4R | A | 8/7/0 | 52015 | 22/21/18 |
| Q96LB1 | MRGX2 | A | 14/14/0 | 9231 | 17/17/17 |
| Q96LA9 | MRGX4 | A | 1/1/0 | 12706 | 19/18/20 |
| **P48039** | MTR1A | A | 6/1/5 | 96729 | 53/21/54 |
| P49286 | MTR1B | A | 4/0/4 | 91932 | 51/19/51 |
| **P25103** | NK1R | A | 11/5/6 | 103865 | 21/20/21 |
| P49146 | NPY2R | A | 1/0/1 | 89832 | 24/26/24 |
| P30989 | NTR1 | A | 2/2/0 | 80910 | 17/17/17 |
| P30559 | OXYR | A | 1/0/1 | 66651 | 16/34/19 |
| Q9Y5Y4 | PD2R2 | A | 3/0/3 | 55542 | 25/25/21 |

| | | | | | |
|---|---|---|---|---|---|
| P43116 | PE2R2 | A | 3/3/0 | 5173 | 13/15/12 |
| P43115 | PE2R3 | A | 2/2/0 | 6001 | 25/22/22 |
| **P35408** | PE2R4 | A | 2/1/1 | 20390 | 11/12/12 |
| P25105 | PTAFR | A | 1/0/0 | 43804 | 23/21/21 |
| Q99500 | S1PR3 | A | 4/4/0 | 40642 | 45/19/43 |
| Q9H228 | S1PR5 | A | 1/1/0 | 27273 | 38/34/38 |
| P21731 | TA2R | A | 2/0/0 | 5300 | 22/29/22 |
| P30518 | V2R | A | 2/2/0 | 39048 | 19/19/14 |
| **Q16602** | CALRL | B1 | 6/4/2 | 22591 | 52/52/23 |
| Q13324 | CRFR2 | B1 | 1/1/0 | 25918 | 29/25/27 |
| Q02643 | GHRHR | B1 | 1/1/0 | 16767 | 33/41/32 |
| P48546 | GIPR | B1 | 1/1/0 | 17173 | 44/45/44 |
| O95838 | GLP2R | B1 | 1/1/0 | 13877 | 32/32/32 |
| P41586 | PACR | B1 | 4/4/0 | 23520 | 29/42/29 |
| **Q03431** | PTH1R | B1 | 4/3/1 | 19756 | 25/24/23 |
| P49190 | PTH2R | B1 | 1/1/0 | 17873 | 44/44/43 |
| P47872 | SCTR | B1 | 2/2/0 | 21881 | 36/45/35 |
| P32241 | VIPR1 | B1 | 1/1/0 | 22749 | 35/44/32 |
| Q86Y34 | AGRG3 | B2 | 2/2/0 | 21765 | 9/11/10 |
| **P41180** | CASR | C | 9/4/5 | 40116 | 21/24/23 |
| Q9UBS5-2 | GABR1 | C | 1/0/1 | 20205 | 15/31/31 |
| **O75899** | GABR2 | C | 8/3/5 | 21450 | 14/14/27 |
| Q5T848 | GP158 | C | 3/0/3 | 7719 | 4/4/5 |
| **Q14416** | GRM2 | C | 9/3/6 | 36733 | 41/44/42 |
| Q14833 | GRM4 | C | 1/1/0 | 37849 | 37/28/42 |
| Q14831-3 | GRM7-3 | C | 1/0/1 | 37310 | 36/28/40 |
| Q9ULV1 | FZD4 | F | 1/0/1 | 13243 | 22/22/40 |
| Q13467 | FZD5 | F | 1/0/1 | 13433 | 22/22/22 |
| O75084 | FZD7 | F | 1/1/0 | 13381 | 22/23/44 |

[1] GPCRs that have been determined in both active and inactive states were highlighted with bold red fonts.

[2] Numbers of experimentally determined GPCR structures in all, active, inactive, and intermediate states.

[3] Maximum template sequence identities from template databases of the standard PDB70 (for the original AlphaFold2 pipeline) and active/inactive/intermediate state GPCR databases (for this work). Templates with high sequence identities (>70%) were excluded from template selection.

**Table S3.** The list of GPCR-ligand complexes for the protein-ligand docking benchmark

| UniProt ID | Gene name | Class | State | Protein-ligand complexes[1] |
|---|---|---|---|---|
| P08908 | 5HT1A | A | Active | 7E2Y+SRO(2,13) |
| P28221 | 5HT1D | A | Active | 7E32+SRO(2,13) |
| P28566 | 5HT1E | A | Active | 7E33+HVU(1,17) |
| P28223 | 5HT2A | A | Active | 6WHA+U0G(7,23) |
| | | | Inactive | 6WH4+89F(2,24), 6WGT+7LD(3,24), 6A94+ZOT(4,22) |
| P08913 | ADA2A | A | Inactive | 6KUY+E39(2,18), 6KUX+E3F(3,25) |
| P18089 | ADA2B | A | Active | 6K41+CZX(2,15) |
| P18825 | ADA2C | A | Inactive | 6KUW+E33(3,25) |
| P08588 | ADRB1 | A | Active | 7BU6+E5E(2,12), 7BTS+ALE(3,13) |
| | | | Inactive | 7BVQ+CAU(6,22) |
| P21728 | DRD1 | A | Active | 7JVQ+OR9(0,20), 7CKW+G3C(1,21), 7CRH+GBU(1,22), 7JV5+SK0(1,20), 7JVP+SK9(1,22), 7LJC+SK0(1,20), 7CKX+G3O(2,24), 7CKZ+LDP(2,11), 7LJD+LDP(2,11) |
| Q6DWJ6 | GP139 | A | Active | 7VUG+7ZQ(5,22), 7VUH+7ZQ(5,22), 7VUI+7ZQ(5,22), 7VUJ+7ZQ(5,22) |
| Q96LB1 | MRGX2 | A | Active | 7S8N+8IU(3,23), 7S8O+8IU(3,23) |
| P48039 | MTR1A | A | Active | 7DB6+JEV(4,19) |
| | | | Inactive | 6ME2+JEV(4,19), 6ME4+ML2(4,18), 6ME5+AWY(4,18), 6ME3+JEY(5,23), 6PS8+JEY(5,23) |
| P49286 | MTR1B | A | Inactive | 6ME9+JEV(4,19), 6ME6+JEY(5,23), 6ME7+JEY(5,23), 6ME8+JEY(5,23) |
| P25103 | NK1R | A | Inactive | 6HLL+GBK(5,22) |
| P43116 | PE2R2 | A | Active | 7CX2+P2E(12,25) |
| P43115 | PE2R3 | A | Active | 6AK3+P2E(12,25) |
| P35408 | PE2R4 | A | Active | 7D7M+P2E(12,25) |
| Q99500 | S1PR3 | A | Active | 7EW4+JF9(4,23) |
| Q14416 | GRM2 | C | Active | 7E9G+HZR(5,24) |

[1] PDB ID and the ligand name. The numbers of rotatable torsion angles and heavy atoms of ligand are shown in the parentheses.

**Algorithm S1** Modification of input MSA features

**def** remove_msa_for_template_aligned_regions($\{\mathbf{f}_{j,i}^{\text{template\_sequence}}\}$, $\{\mathbf{f}_{k,i}^{\text{msa}}\}\{\mathbf{f}_{k,i}^{\text{deletion\_matrix\_int}}\}$):

**for all** $i \in [1, \ldots, N_{residue}]$ **do**
### determine whether residue $i$ is aligned to any selected templates
    $aligned :=$ **False**
    **for all** $j \in [1, \ldots, N_{template}]$ **do**
        $s_{j,i} := \mathbf{f}_{j,i}^{\text{template\_sequence}}$
        **if** $\mathbf{s}_{j,i} \neq "-"$ **then**
            $aligned :=$ **True**
        **end if**
    **end for**
### modify input MSA features for the residue
    **if** $aligned =$ **True then**
        **for all** $k \in [1, \ldots, N_{sequence}]$ **do**
            $\mathbf{f}_{k,i}^{\text{msa}} := 21$ # 21 for gap
            $\mathbf{f}_{k,i}^{\text{deletion\_matrix\_int}} := 0$
        **end for**
    **end if**
**end for**
**return** $\{\mathbf{f}_{k,i}^{\text{msa}}\}$, $\{\mathbf{f}_{k,i}^{\text{deletion\_matrix\_int}}\}$
